# Deep lipidomic profiling reveals sex dimorphism of lipid metabolism in fibro-calcific aortic valve disease

**DOI:** 10.1101/2024.08.07.606946

**Authors:** Patricia Prabutzki, Michele Wölk, Julia Böttner, Zhixu Ni, Sarah Werner, Holger Thiele, Jürgen Schiller, Petra Büttner, Florian Schlotter, Maria Fedorova

**Affiliations:** Leipzig University, Faculty of Medicine, Institute for Medical Physics and Biophysics, Leipzig, Germany; Center of Membrane Biochemistry and Lipid Research, University Hospital and Faculty of Medicine Carl Gustav Carus of TU Dresden, Dresden, Germany; Heart Center Leipzig at Leipzig University, Department of Internal Medicine/Cardiology, Leipzig, Germany; Department of Cardiology, University Medical Center of the Johannes Gutenberg University Mainz and German Center for Cardiovascular Research - Partner Site Rhine-Main, Mainz, Germany

**Author notes:** Authors contributed equally to this work (Patricia Prabutzki, Michele Wölk). Florian Schlotter, Department of Cardiology, University Medical Center of the Johannes Gutenberg University Mainz and German Center for Cardiovascular Research - Partner Site Rhine-Main, Mainz, Germany;, Maria Fedorova, Lipid Metabolism: Analysis and Integration, Center of Membrane Biochemistry and Lipid Research, University Hospital Carl Gustav Carus and Faculty of Medicine of TU Dresden, Dresden, Germany.

**Keywords:** aortic stenosis, lipidomics, fibro-calcific aortic valve disease, fibrosis, calcification

## Abstract

Fibro-calcific aortic valve disease (FCAVD) is the most common valvular heart disease manifesting in pathological fibro-calcific remodeling of the aortic valve (AV) leaflets, ultimately leading to aortic stenosis. Although lipid dysmetabolism is a driver of FCAVD pathogenesis, the molecular details of the AV lipidome remodeling upon fibrosis and calcification remain largely unknown. Here, we employed advanced lipidomics technologies for deep quantitative profiling of metabolic trajectories in human tricuspid and bicuspid AVs at different pathological stages. Specific extrinsic and intrinsic lipid trends, accompanying the development of fibrosis and calcification, were identified. Importantly, significant differences in lipid signatures between male and female individuals were demonstrated and were attributable to altered sphingolipid metabolism. Taken together, deep lipidomics profiling allowed to identify major molecular events and revealed a high extent of sex-dimorphism in lipidomics signatures of human FCAVD.

## Main

Aortic stenosis (AS) is a severe condition, which is characterized by thickening of the aortic valve (AV) leaflets and fibro-calcific remodeling of the tissue, ultimately leading to the obstruction of the cardiac outflow^1^. Due to its reorganization during the FCAVD progression, tissue section in the non- or mildly diseased as well as fibrotic (progressed) and calcific (end stage) stage can be present in the same AV leaflet. Tissue fibrosis and calcification result in narrowing of the AV opening and induce left ventricular hypertrophy, which consequently manifests in reduced cardiac output and heart failure^2^. Fibro-calcific aortic valve disease (FCAVD) is the most common valvular heart disease in the aging population^3^. Surgical aortic valve replacement (SAVR) or transcatheter aortic valve implantation (TAVI) remain the only treatment options and pose a major economic burden on the health care systems, as a pharmacological therapy does not yet exist.

Identification of pharmacological targets is further complicated by the heterogeneity of the human population. Individuals with congenital AV abnormalities such as uni- and bicuspid AVs typically show earlier onset of FCAVD, attributed to increased mechanical stress, and manifest clinically relevant determinants of symptomatic AS on average 10 years earlier than patients with the physiological tricuspid AV (TAV)^4^. Furthermore, recent evidence revealed sex-related differences in AS with more fibrotic phenotypes present in females as opposed to predominantly calcific remodeling in males^5^. Despite major advances in the field, the cardiovascular scientific research community has yet to clarify the precise molecular mechanisms that govern FCAVD onset and progression to facilitate an effective pharmacological intervention.

Major FCAVD hallmarks include differentiation of valve interstitial cells (VICs) to pro-fibrotic and calcific phenotypes, chronic inflammation, immune cell infiltration and lipoprotein accumulation^6^. Indeed, numerous clinical studies have shown a strong correlation between blood plasma lipid profiles of patients with FCAVD with disease incidence and progression^7–10^. This allowed to identify several lipid-related traits in FCAVD driven by a pathological deposition of low-density lipoprotein (LDL) and lipoprotein(a) (Lp(a)), including the lipoprotein-associated phospholipase A_2_ (LpPLA_2_) mediated conversion of glycerophosphatidylcholine (PC) lipids to the corresponding lyso-forms (LPC), which in turn can be metabolized to bioactive lysophosphatidic acids (LPA) by the action of the secreted phospholipase D, autotaxin^11–13^. Similarly, sphingomyelins (SM) delivered by LDL and Lp(a) are converted to the corresponding ceramides (Cer) by the action of sphingomyelinase (SMase), secreted by infiltrated immune cells or resident valve endothelial cells (VEC)^14^. Such metabolic processing of accumulating lipoproteins promotes further lipid deposition, as the generated LPC significantly increases LDL aggregation rates, whereas Cer are positively correlated with both, LDL particle fusion and aggregation^15^. Furthermore, oxidized LDL and Lp(a) have been shown to promote osteoblastic VIC differentiation and hydroxyapatite mineralized matrix deposition^16,17^. However, pharmacological, lipid-lowering therapy with the highly successful anti-atherosclerotic medication class of HMG-CoA reductase inhibitors (statins) failed to show clinical benefits in FCAVD^18–20^.

Even though different lipid-related traits of FCAVD progression have been observed, changes in the lipid composition of the affected AV tissues have not been comprehensively characterized. Indeed, qualitative and quantitative descriptions of the specialized lipidomes of human tissues under physiological and pathological conditions have been only sporadically addressed^21–23^. Here, we provide a systematic qualitative and semi-quantitative molecular description of the human AV lipidome covering mildly diseased, fibrotic and calcific phenotypes. In-depth lipidomics profiling revealed metabolic trajectories associated with the development of fibrosis and calcification both in tri- and bicuspid human AVs. Importantly, we identified significant sex-dimorphism in FCAVD development at the molecular level, which was mainly attributed to sphingolipid metabolism. Finally, all raw and processed lipidomics data produced in this study are made publicly available via Metabolomics Workbench^24^ resource, representing the first comprehensive human AV reference lipidome available for the community for further analysis and data interrogation^25^.

## Results

### Reference lipidome of the human AV

To define the molecular composition of the human AV lipidome, we first performed deep lipidomics profiling of pooled samples representing mildly diseased sections of dissected tricuspid AVs (TAV) of patients undergoing SAVR for severe AS. Mildly diseased TAV regions of 23 patients (13 males and 10 females) with an average age of 72 years were used for lipid extraction (Supplementary Data Table 1). Reverse phase chromatography (RPC) coupled online with mass spectrometry (MS) analysis performed in an untargeted and targeted manner (see Methods section for details) allowed to identify 1073 lipid molecular species (Supplementary Data Table 2), of which 480 lipids were semi-absolutely quantified (Supplementary Data Table 3 and Fig. 1a). The human TAV lipidome is represented by 28 lipid subclasses spanning over 6 orders of magnitude in lipid concentrations with cholesteryl ester (CE) 18:2 (5.9 nmol/mg wet tissue weight) being the most and sphingomyelin (SM) 32:0;O3 (0.003 pmol/mg) being the least abundant species. The most abundant lipid classes in human TAV lipidome were CE (total amount 10.80 nmol/mg), followed by triacylglycerides (TG; 1.65 nmol/mg), PC (0.8 nmol/mg), SM (0.76 nmol/mg), and ether glycerophosphatidylethanolamines (etherPE; 0.27 nmol/mg).

**Fig. 1.**
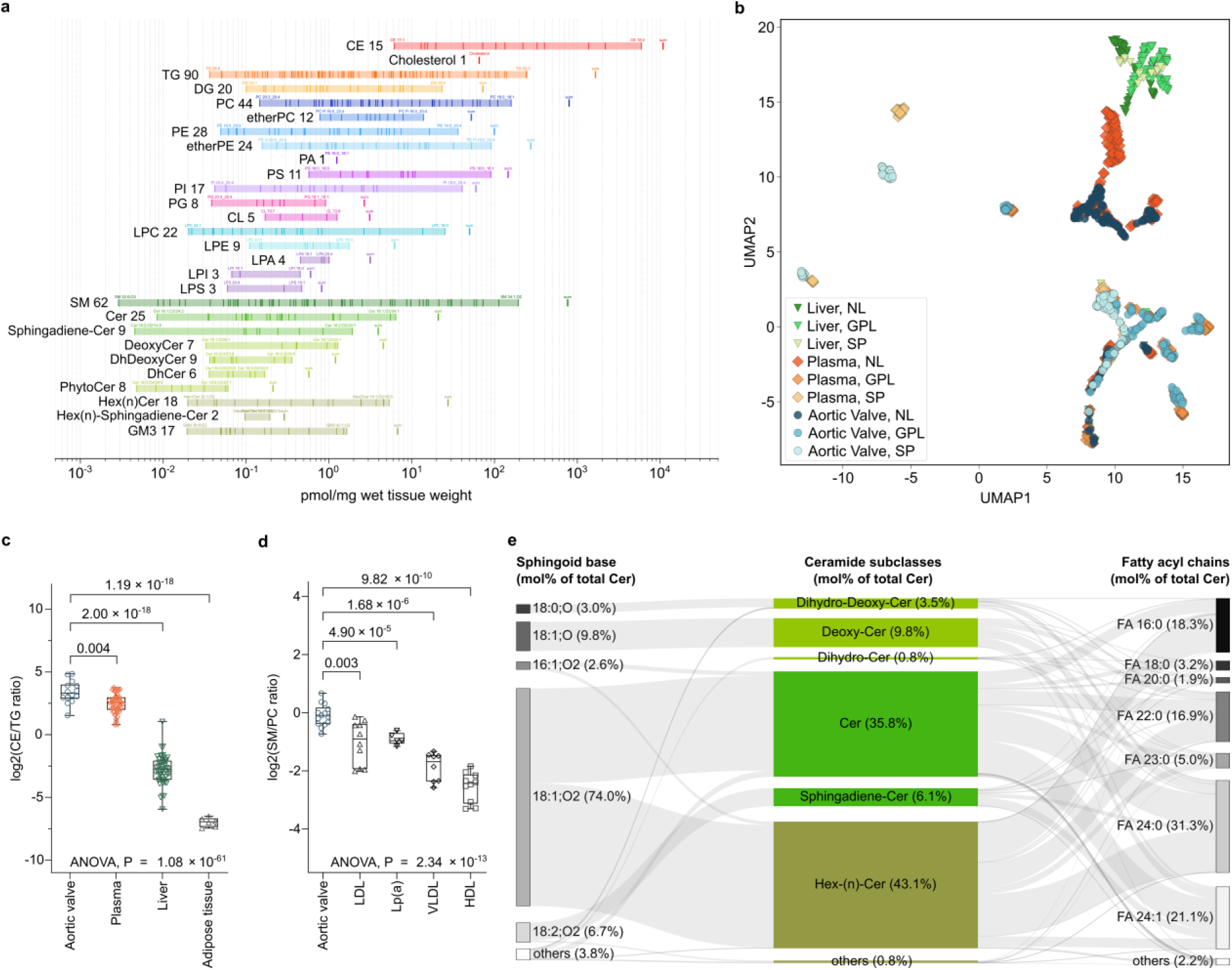
Deep lipidomics profiling of mildly diseased human TAV. **a,** Quantitative distribution of lipid subclasses and corresponding molecular species in mildly diseased human TAV of elderly individuals. Total concentration for each lipid subclass is shown by the bold line (SUM) and the concentrations of each lipid molecular species are marked by thin lines within shaded region. **b,** Topological comparison using uniform manifold approximation and projection (UMAP) of human TAV, blood plasma and liver lipidomes. **c,** Total CE to TG ratio in TAV (n=14), blood plasma (n=50), liver (n=49) and adipose tissue (n=6) lipidomes. **d,** Total SM to PC ratio in TAV (n=14) and different lipoprotein fractions (n=5). P values were calculated by Welch’s T-test or one-way ANOVA across all groups. Boxplot elements represent centerline, mean; box limits, 25% and 75% quartiles; vertical lines connect minimum and maximum values. Dots represent independent samples. **e,** Sankey plot illustrating the relative abundance of TAV ceramide (Cer) subclasses (central), composed of the corresponding sphingoid bases and fatty acyl chains shown on the left and right sides, respectively.

Considering the histological organization of AVs, high enrichment in neutral lipids (NL) could be attributed to the accumulation of blood lipoproteins in mildly diseased AV sections of elderly individuals. To trace the source of lipid infiltration at the molecular level, we compared the TAV lipid composition with previously published lipidomes of human blood plasma, as a source of circulating lipids^26^, and liver, as a major lipoprotein producing organ^27^. To this end, we employed uniform manifold approximation and projection (UMAP) algorithm for dimensionality reduction and topological analysis of tissue lipidome similarities. Z-scored lipid abundances and structural features were used as parameters for the projections. UMAP clearly illustrated the close similarity between TAV and plasma lipidomes, whereas liver lipids presented a different topology (Fig. 1b). Specifically, we observed a very close overlap between the TAV and plasma CE with all species sharing similar topology (Extended Data Fig. 1a). TG lipids formed several topological clusters (Extended Data Fig. 1b), with TAV TGs distributed between two clusters - one shared between all three biological samples (mostly TG with 50-54 carbon atoms), and the second formed by TAV and blood plasma TGs. The same was true for PC and LPC lipids (Extended Data Fig. 1c), whereas sphingolipids (SP) showed tissue-specific topologies except for SM, which closely overlapped between TAV and plasma lipidomes (Extended Data Fig. 1d). Taken together, with the exception for ceramide (Cer) lipids, the topological analysis confirmed the close similarity of the TAV and blood plasma lipidomes, especially in respect to the major components of circulating lipoproteins (CE, TG, PC and SM). Furthermore, when compared in terms of total CE to TG ratio (Fig. 1c), the TAV lipidome of mildly diseases sections, was more similar to blood plasma^26^, than other NL-rich tissues including liver^27^ or adipose tissue^26^.

Next, we examined whether lipid molecular signatures are useful to define which circulating lipoproteins contribute to the lipidome of mildly diseased TAV. Based on immunohistochemistry and proteomics data, a major contribution of both LDL and Lp(a) in TAV lipid deposition is proposed^28,29^. Indeed, based on the CE to TG ratio a more significant impact of CE-rich rather than TG-rich lipoproteins was confirmed (Fig. 1c). Additionally, we compared the ratio of SM to PC lipids in TAV versus LDL, Lp(a), very low-density (VLDL) and high-density (HDL) lipoproteins for which different relative contributions of SM and PC lipids were reported^30,31^. Comparison of the TAV specific SM/PC ratio to the previously published data for various lipoprotein fractions^32–35^ revealed its close similarity to LDL and Lp(a) (Fig. 1d). This comparative analysis implies that a substantial fraction of the TAV lipidome, especially with respect to NL and sphingolipids (SP), derives from retained and accumulated LDL and Lp(a) particles in mildly diseased TAV regions of elderly individuals. This is further supported by the fact that CE 18:2, a marker of extracellular LDL accumulation^23^, is by far the most abundant lipid in mildly diseased TAV (Fig. 1a).

Interestingly, the lipidome of mildly diseased TAV revealed a remarkable richness and diversity of SP lipids, when compared to blood plasma. In addition to highly abundant SM sharing plasma SM topology, various subclasses of Cer formed TAV-specific clusters (Extended Data Fig. 1d). It is particularly interesting since these lipids recently gained high significance as potential markers for cardiovascular diseases^36^. TAV Cer showed very high structural diversity, both in terms of sphingoid bases and fatty acyl (FA) chains (Fig. 1e), with sphingosine (18:1;O2) being the most abundant base (74% of all Cer), followed by deoxysphingosine (18:1;O; 9.8%) and sphingodienine (18:2;O2; 6.7%), whereas the most abundant acyl chains were represented by FA 24:0 (31.3%), FA 24:1 (21.1%), and FA 16:0 (18.3%). Furthermore, the total amount of glycosphingolipids represented by hexosylceramides (Hex_n_Cer) and gangliosides (GM3) (34.6 pmol/mg), even exceeded that of Cer (24.9 pmol/mg) in the TAV tissue (Fig. 1a). Despite their high abundance, Hex_n_Cer were structurally less heterogeneous and almost entirely composed of sphingosine backbones and FA 16:0, FA 22:0 or FA 24:0 acyl chains.

Taken together, using deep lipidomics profiling and semi-absolute quantification, we defined the reference lipidome of mildly diseased human TAV of patients with FCAVD. By comparing the TAV lipidome with previously published lipidomes of other human tissues (adipose tissue, liver, and blood plasma), we could demonstrate a strong similarity between TAV and plasma lipid compositions. Furthermore, using the CE/TG and SM/PC ratio as a proxy for different lipoproteins, enrichment of LDL and Lp(a) particles over VLDL and HDL was confirmed. Interestingly, in addition to the expected accumulation of typical lipoprotein-derived lipids (CE, TG, PC and SM), we could demonstrate a large diversity of ceramides as a specific molecular feature of the human TAV lipidome.

### FCAVD metabolic trajectories revealed by TAV fibrotic and calcific lipidomes

Next, we focused on identifying the molecular signatures of the TAV lipidome in the pathological stages of FCAVD. To this end, the AV of patients undergoing valve replacement for severe AS were dissected into different regions, namely mildly diseased, fibrotic and calcific sections (Fig. 2a). Relative to the mildly diseased sections, total lipid content increases drastically in fibrotic (2.3-fold) and calcific (2-fold) tissue sections. CE, cholesterol and SPs contributed the most to the elevated lipid load. Thus, CE concentrations in fibrotic and calcific sections were increased 2.7- and 2.3-fold, respectively (Fig. 2b). Free cholesterol was significantly elevated in the fibrotic (2.6-fold) and even further increased (3.0-fold) in the calcific state (Fig. 2c). Interestingly, total TG load did not change much between different FCAVD progression stages and in general showed high inter-individual variability (Fig. 2c). Similarly, no increase in the load of glycerophospholipids (GPL) was observed, whereas SPs showed large accumulation in fibrotic (2.0-fold) and calcific (1.9-fold) sections of TAV (Fig. 2d). Quantitative comparison of all lipid subclasses between mildly diseased, fibrotic and calcific TAV sections is provided in Extended Data Fig. 2.

**Fig. 2.**
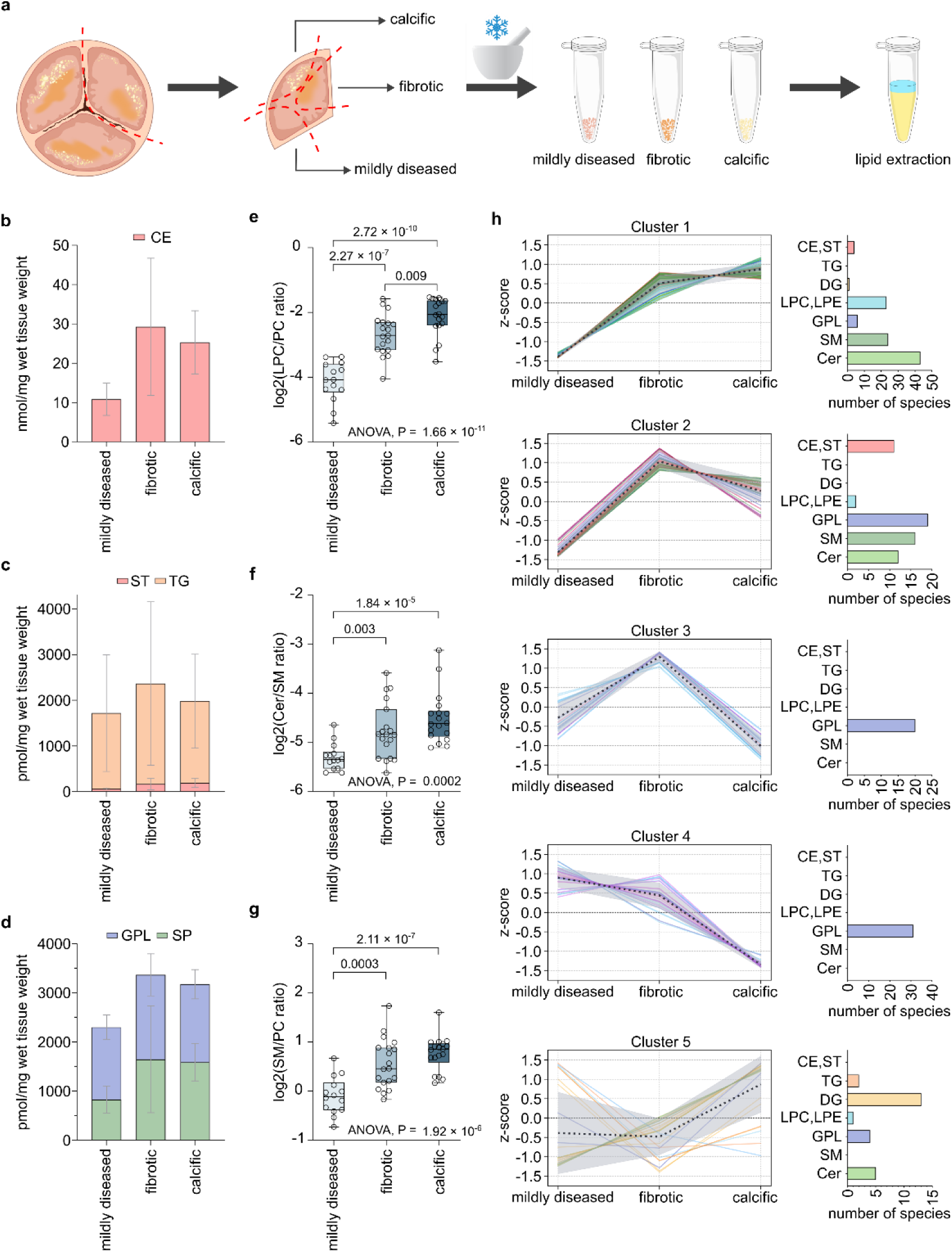
Aortic valve lipidome remodeling in FCAVD of TAV. **a,** Schematic representation of AV tissue dissection into mildly diseased, fibrotic and calcific sections for a separate lipid extraction. **b-d,** Bar plots showing the mean total concentration of cholesteryl esters (CE) (**b**), triacylglycerides (TG) and free cholesterol (ST) (**c**), glycerophospholipids (GPL) and sphingolipids (SP) (**d**) in mildly diseased, fibrotic and calcific tissue sections. Margins indicate standard deviation. **e-g,** Total LPC to PC (**e**), Cer to SM (**f**), and SM to PC (**g**) ratios in mildly diseased, fibrotic and calcific TAV sections. P values were calculated by T-test or one-way ANOVA across all groups. Boxplot elements represent centerline, mean; box limits, 25% and 75% quartiles; vertical lines connect minimum and maximum values. Dots represent biologically independent samples. Color-coding indicates pathophysiological stage: light blue – mildly diseased; blue - fibrotic; dark blue - calcific. **h,** Lipid Trend Analysis via Gaussian mixture model clustering (number of clusters = 5) of z-scored concentrations for 237 significantly regulated lipids (determined by ANOVA P < 0.05) along FCAVD progression in TAV. Line color indicated different lipid classes: red - cholesterol and CE; orange - TG; yellow - DG; LPC and LPE - light blue; GPL - dark blue; SM - dark green; Cer - light green.

To investigate the pathophysiological lipidome alterations in TAV further, we focused on the known lipid-related traits of FCAVD progression, namely (I) the LDL – LpPLA_2_ – LPC – LPA axis, and (II) SMase induced Cer accumulation. As expected, validating our approach, we observed a significant increase in the absolute LPC concentrations (Supplementary Data Table 3) as well as the ratio of total LPC to PC lipids in the fibrotic (2.8-fold) and calcific (4.0-fold) TAV relative to the mildly diseased sections (Fig. 2e), indicating high LpPLA_2_ activity promoting consequent lipoprotein aggregation and deposition. Similar trends, (1.7- and 1.8-fold increase in fibrotic and calcific sections, respectively), were observed regarding the LPE to PE ratio (Extended Data Fig. 3).

Several reports suggested that the fibro-calcific response in FCAVD is not driven by LPC itself, but rather by LPC-derived LPA produced by the phospholipase D autotaxin^12^. Despite previously published high LPA concentrations in AV tissue^21,37^, we were unable to detect any endogenous LPA or PA species in lipid extracts obtained by conventional lipid extraction, i.e., Folch method (Extended Data Fig. 4). Considering the proposed relevance of LPA in FCAVD pathology, we therefore employed acidified methanol-chloroform based extraction^38^, followed by targeted LC-MS/MS analysis of FCAVD tissues in order to analyze PA and LPA specifically (Extended Data Fig. 4). To note, PA and LPA analysis requires careful analytical optimization, as both extraction and MS analysis in complex biological matrices can result in artificial PA and LPA generation either by acid-mediated hydrolysis or in-source fragmentation of other GPL^39–41^. Considering all the analytical limitations mentioned above, we detected and quantified one PA and four LPA molecular species, of which only LPA 16:0 showed significant difference and was slightly elevated in calcific TAV sections (Extended Data Fig. 4 and Supplementary Data Table 3).

Next, we investigated the ratio between Cer and SM lipids as a second prominent trait associated with increased lipoprotein deposition^15^. Not only the total SM content was 1.9 and 1.8 fold higher in fibrotic and calcific TAV relative to mildly diseased sections, but also the Cer/SM ratio rose 1.5- and 1.8-fold, respectively with FCAVD progression. This indicates that total Cer content increase was even more pronounced due to the conversion of infiltrating SM to the corresponding Cer by the action of SMase, priming lipoprotein aggregation and deposition (Fig. 2f). Generally, LDL aggregation was shown to correlate positively with SM and Cer lipids and negatively with PC lipids^31^. We could further confirm this lipid-related FCAVD trait and show that the total SM to PC ratio steadily increased with disease progression corresponding to 1.6- and 1.8-fold higher values in fibrotic and calcific sections relative to mildly diseases TAV (Fig. 2g). Thus, by defining mildly diseased, fibrotic and calcific TAV lipidomes, we could confirm, in a quantitative manner, the major traits in FCAVD pathogenesis associated with increased LDL and Lp(a) aggregation and deposition, namely PC to LPC and SM to Cer conversions, serving as a proxy for LpPLA_2_ and SMase activities.

Lipid trends described above represent an important part of the TAV lipidome remodelling driven mainly by lipoprotein aggregation and deposition. However, during FCAVD pathogenesis, the TAV lipidome undergoes alterations induced not only by lipoprotein infiltration but also due to the intrinsic remodelling driven by VIC transition to myofibroblastic and osteoblastic phenotypes, VEC endothelial to mesenchymal transition and immune cell infiltration. To dissect extrinsic and intrinsic lipidomic signatures of the TAV pathology, we performed Lipid Trends Analysis, utilizing Gaussian mixture model clustering of 237 lipid species significantly regulated between mildly diseases, fibrotic and calcific TAV sections (ANOVA p < 0.05); Fig. 2h). The first two clusters can be attributed to the extrinsic trends mediated by lipoprotein infiltration characterized by progressive (cluster 1) or fibrotic stage tipping (cluster 2) lipid depositions. Thus, clusters 1 and 2 are formed by 101 and 60 lipids, respectively, and include major lipid species likely originating from lipoprotein accumulation - cholesterol, CE, and SM. Interestingly, most of the CE belonged to cluster 2 showing that deposition of lipoproteins occurs mainly in the fibrotic stage, and the total load of CE lipids does not increase during the progression from the fibrotic to the calcific stage. On the other hand, cluster 1, in addition to the infiltrated lipids (cholesterol and SM) also included LPC, LPE and Cer, produced locally from lipoprotein-derived lipids by the action of LpLPA_2_ and SMase. These lipids showed progressive accumulation over the pathological stages of FCAVD with steep elevation already at the fibrotic stage and moderate increase in calcific TAV. Thus, cluster 1 and 2 can be assigned as “lipoprotein infiltration” and “lipoprotein processing” trends, respectively. Interestingly, cluster 1 included 13 glycoSP namely species carrying one, two or three hexose moieties. Hex_n_Cer can derive both from infiltrating lipoproteins directly and be produced locally from Cer lipids. In the context of atherosclerosis, it was shown that Hex_n_Cer can promote cholesterol accumulation by increasing LDL uptake and inhibiting HDL-mediated cholesterol efflux in macrophages, inducing foam cells formation, increase monocyte adhesion and stimulate vascular smooth muscle cells proliferation^42^. However, in FCAVD the role of Hex_n_Cer remains undefined.

Cluster 3 and 4 showed not only different trends but also very different compositions compared to the lipoprotein-associated clusters 1 and 2, and thus, we propose that they might represent TAV intrinsic lipidome remodeling. Indeed, neither CE nor SM species are present in these clusters, which are solely represented by various GPL lipids (Fig. 2h). Abundances of lipids from cluster 3 tip at fibrotic stage and rapidly decline with TAV calcification, possibly indicating the increase in TAV cell populations due to VIC and VEC transformations and immune cell influx alone with fibrosis development. Indeed, it contains 20 lipids, 18 of which are represented by major membrane building PC, PE and etherPE species. Cluster 4, on the other hand, shows progressive decline in abundances of 31 lipid species, especially evident for calcific TAV. In addition to a few PC, PE, and PUFA-rich etherPE lipids, cluster 4 includes cardiolipins (CL; 4 out 5 detected species) and phosphatidylserine (PS; 10 out of 11 detected species) lipids. We attributed cluster 4 to the major tissue degenerative events occurring in calcific TAV accompanied by the massive loss of PUFA etherPE due to increased oxidative stress and cell death, evidenced by a sharp decline of mitochondria-specific CL lipids. PS lipids depletion in calcific TAV represents an interesting phenomenon and serves as a specific marker of progressive tissue calcification. Finally, the remaining cluster 5 combines a set of mixed lipids species (25 lipids) with no common trend. A large portion of the lipids in this cluster are diacylglycerols (DG) showing largely diverse trends based on their acyl chains compositions. Thus, PUFA DG increased with FCAVD pathology and DG with saturated acyl chains showed an opposite trend.

Taken together, using quantitative lipidomics of FCAVD disease stages, we showed that TAVs accumulate large amounts of lipids mostly at the fibrotic stage. CE provide the major impact in this lipid build-up, followed by free cholesterol and SP lipids, in agreement with the concept of pathological LDL and Lp(a) deposition. Importantly, using Lipid Trends Analysis, we could dissect extrinsic (associated with lipoprotein infiltration) and intrinsic (driven by TAV cell populations) metabolic trajectories of lipidome remodeling in FCAVD progression. Thus, we demonstrated that lipoprotein-derived lipids undergo pathological processing in TAV leading to the accumulation of lysoGPL (LPC and LPE) and Cer, mediated by the activity of LpPLA_2_ and SMase, respectively. Additionally, glycosphingolipids (Hex_n_Cer) showed progressive accumulation over the disease stages. Among intrinsic pathways, lipidome signatures indicated increased cell density at the fibrotic stage followed by tissue degeneration (PUFA-etherPE and CL depletion) in calcific TAV sections. Interestingly, one of the most evident hallmarks of the calcific TAV lipidome was a massive reduction in PS lipids, which are probably involved in the calcification process due to their anionic nature and, thus, can serve as markers of FCAVD progression.

### Sex dimorphism in lipidome remodeling of FCAVD

Multiple lines of evidence indicate that biological sex plays an important role in FCAVD progression, with women developing a more fibrotic phenotype with the same hemodynamically defined degree of AS, while men are characterized by a higher extent of calcification^43–45^. Indeed, AV from female patients included in this study generally presented with more of a fibrotic phenotype in comparison to the samples of male patients, which showed higher calcification (Fig. 3a). Thus, we decided to compare the differences in lipidome signatures of AV sections for male and female patients.

**Fig. 3.**
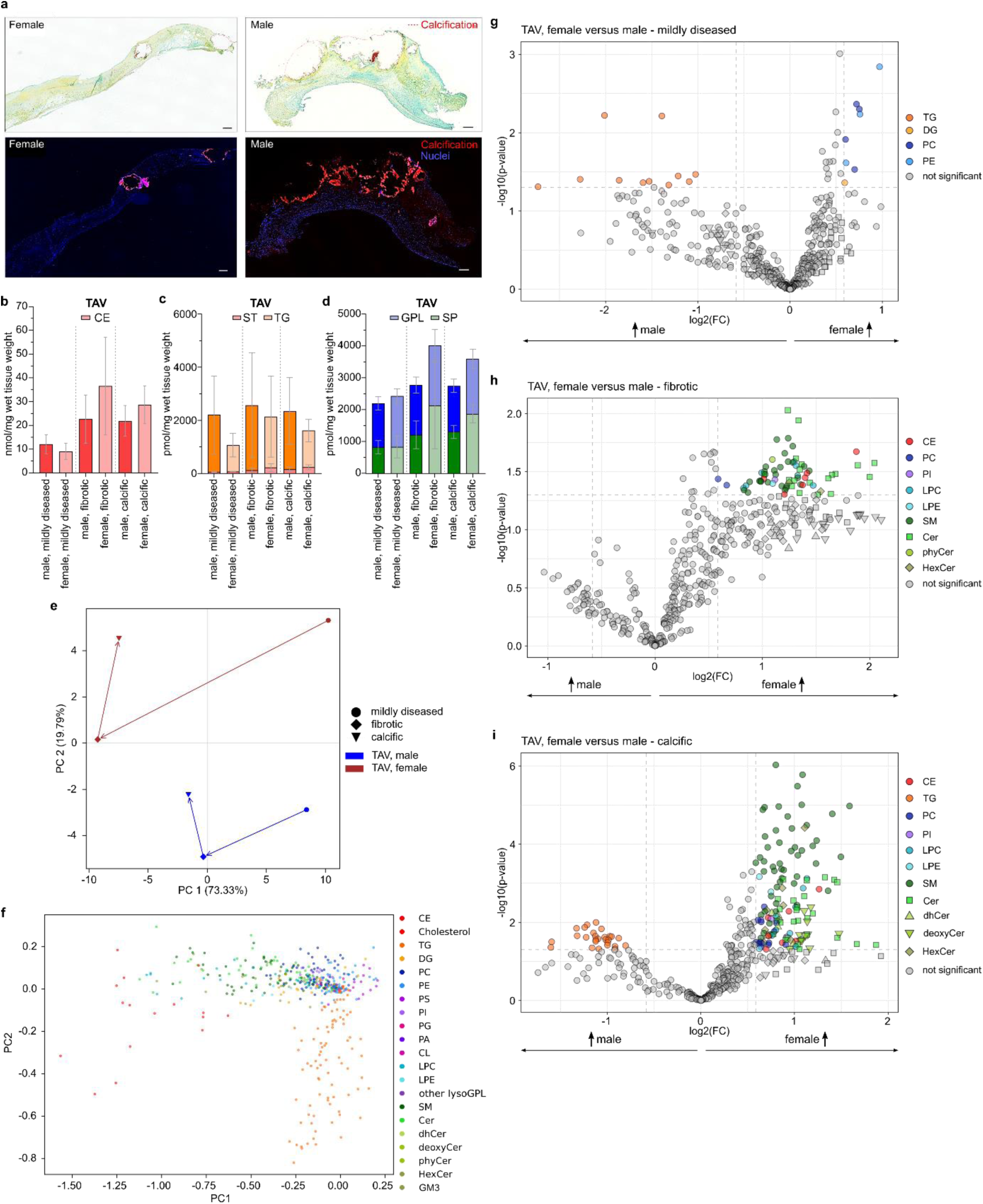
Sex dimorphism in lipidome remodeling in FCAVD. **a,** Extracellular matrix remodeling in female and male FCAVD TAV sections visualized with Movat‘s pentachrome staining (top panels) shows collagen (yellow) and glycosaminoglycans (turquoise) and OsteoSense and DAPI staining (bottom panels) for calcific mineralization (red) and nuclei (blue), respectively. Bar correspond to 100 µm. **b-d,** Bar plots showing the mean total concentration of cholesteryl esters (CE) (**b**), triacylglycerides (TG) and free cholesterol (ST) (**c**), glycerophospholipids (GPL) and sphingolipids (SP) (**d**) in mildly diseased, fibrotic and calcific TAV tissue sections of male and female patients, respectively. Margins indicate standard deviation **e-f,** Scoring (**e**) and loading (**f**) plots for Lipidome Trajectories principal component analysis (PCA) of TAV lipidomes of female and male individuals across different pathophysiological stages of FCAVD. Each marker represents an individual lipid species colored by the corresponding lipid subclass **g-i,** Volcano plots illustrating lipid species significantly regulated between female versus male individuals in mildly diseased (**g**), fibrotic (**h**) and calcific (**i**) TAV tissue sections determined by fold change (FC > 1.5) analysis and T-test (P < 0.05).

First, striking differences in the extent of the total lipid deposition between male and female TAVs were detected (Figs. 3b-d). Mildly diseased TAV tissues of males had a higher lipid load (16.5 vs 12.6 nmol/mg wet tissue weight in male and female TAV respectively), mainly due to the elevated content of total CE and TG lipids. However, upon development of fibrosis and calcification female TAV sections accumulated higher amounts of lipids. Thus, relative to mildly diseased sections, male TAV increased their lipid load up to 1.7 and 1.6-fold, whereas female patients showed a much higher lipid deposition with a 3.4- and 2.7-fold increase in fibrotic and calcific stages, respectively. These differences occurred mostly at the fibrotic stage and were mainly driven by the 4-fold increase in CE lipids in female diseased TAVs (Fig. 3b). Overall, all lipid classes except GPL showed higher fold changes in both fibrotic and calcific stages of female patients (Figs. 3c and d) when compared to male TAV sections.

To have a deeper look into sex-dimorphic FCAVD lipid phenotypes, we performed Lipidome Trajectories principal component analysis (PCA) to track lipidome remodeling at different pathological states in a sex-specific manner (Figs. 3e and f). The first principal component (PC1), which explained over 73% of the variables, presented the major FCAVD trends discussed above – accumulation of CE (red), lysoGPL (light blue) and SP (green) lipids (Fig. 3f). The trajectory along PC1 further illustrates that female patients accumulate much higher loads of lipoprotein-derived lipids along the FCAVD progression. PC2, which explained around 20% of lipid-derived variables, illustrated the major sex-specific differences between the pathological stages, which seems to be driven mainly by CE, SP, and TG lipids (orange) (Fig. 3f). Indeed, TG lipids, which did not show any significant differences when data from male and female patients were analyzed together, here, emerge as one of the main sex-specific trends already in the mildly diseased stage.

Next, we performed a pairwise comparison of female vs male TAV lipidomes at each pathological stage (Figs. 3g-i). Thus, male mildly diseased TAV contained higher amounts of TG lipids carrying PUFA acyl chains (C56-C62, with 5 to 9 double bounds), whereas female TAV had higher levels of PUFA containing plasmalogen PE (P-PE) and PC carrying mostly saturated and monounsaturated acyl chains (MUFA) (Fig. 3g). However, the major differences became apparent when comparing fibrotic and calcific sections, in which 66 and 161 lipids were differentially regulated in female vs male fibrotic and calcific TAV sections, respectively (Figs. 3h and i). In fibrotic TAV sections, all significantly different lipids were upregulated in female patients and included CE, SM, Cer, and lysoGPL lipids, indicating a higher pathological conversion (PC to LPC and SM to Cer) of infiltrating lipids, which in turn results in a higher lipoprotein aggregation and deposition.

This sex-driven dimorphism in lipid accumulation was further evident in calcific sections (Fig. 3i). Here, 134 and 27 lipids were significantly upregulated in female or male calcific TAV sections, respectively. The 27 lipids, significantly upregulated in male TAV were all TG lipids, among which PUFA-rich species dominated again. Interestingly, at the quantitative level, the overall TG load in calcific male TAV tissue did not increase relative to mildly diseased sections (roughly 2.2 nmol/mg wet tissue weight in both mildly diseased and calcific male TAV), whereas female TAV increased their TG load 1.4-fold (1.0 vs 1.4 nmol/mg wet tissue weight in mildly diseased and calcific female TAV, respectively) (Fig. 3c). Thus, significantly higher concentrations of particular TG in male vs female patients indicate sex-specific differences at the level of specific molecular species, namely PUFA-rich TG, in calcific male TAV. Similar to the fibrotic stage, major lipid classes upregulated in calcific female TAV are represented by CE, lysoGPL and SP lipids (Fig. 3i).

Almost all measured SP subclasses were upregulated in female calcific TAV including SM and regular Cer, as well as dhCer, deoxyCer, sphingadienine-Cer, phytoCer, and glycosylated species (Fig. 3i). While SM were represented by a diverse composition of molecular species ranging from SM(28:1;O2) to SM(54:2;O3), accumulated Cer showed distinct acyl chains enrichment. Independent of the sphingoid base, the vast majority of Cer lipids, which were found to be accumulating in female calcific TAV sections, carried very long chain fatty acids (VLCFA) ranging from C22 to C26. Thus, we propose that the SP pool in calcific TAV of female patients can be formed not only due to the conversions of lipoprotein derived SM to the corresponding Cer but also via AV-local SP metabolism. In favor of this hypothesis, we could detect sex-specific upregulation of dhCer, which are precursors of Cer lipids in *de novo* synthesis pathways, carrying the same VLCFA acyl residues as elevated Cer species.

Taken together, we demonstrated significant sex dimorphism in lipidome signatures of FCAVD. Thus, mildly diseased TAV sections of male patients had higher lipid contents due to the elevated CE and TG lipids. However, female TAV accumulated much more lipids with the development of fibrosis, mainly due to the massive deposition of lipoprotein-derived lipids and their metabolic conversion. SP lipids emerged as a main discriminator of sex-specific lipidomics remodeling in FCAVD, with female individuals accumulating significantly higher levels of SM and Cer lipids.

### Lipidomics signatures discriminate tricuspid and bicuspid valve morphology

Finally, we compared lipidomics signatures of TAV vs BAV. Overall, we did not detect major differences in BAV lipidomes when compared to TAV lipidomes of the corresponding FCAVD progression stages. Pairwise comparison between pathological stages of BAV and TAV, revealed that mildly diseased sections of BAV contained more SP lipids represented specifically by VLCFA species of a variety of SP subclasses (phytoCer, Cer, sphingadienine-Cer, SM, GM3 and Hex2Cer) (Fig. 4a). Fibrotic and calcific BAV showed lower contents of TG lipids (Figs. 4b and c). Interestingly, in calcific BAV downregulated lipids were PUFA-rich TG as well as VLCFA deoxyCer.

**Fig. 4.**
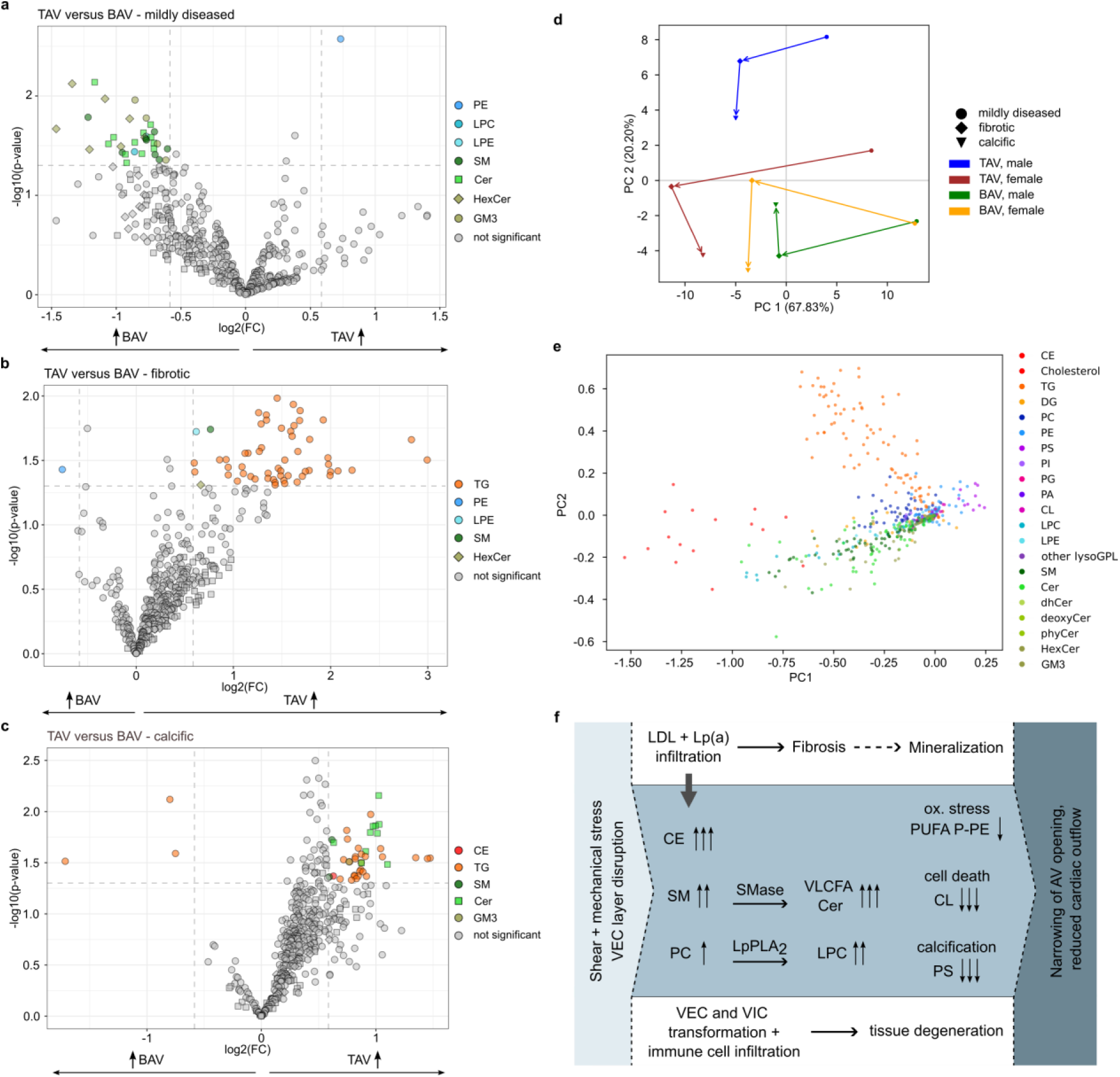
Lipidomics signatures discriminate tricuspid (TAV) from bicuspid (BAV) AV in FCAVD. **a-c,** Volcano plots illustrating lipid species significantly regulated between TAV versus BAV lipidomes of mildly diseased (**a**), fibrotic (**b**) and calcific (**c**) AV tissue sections determined by fold change (FC > 1.5) analysis and T-test (P < 0.05). **d-e,** Scoring (**d**) and loading (**e**) plots for Lipidome Trajectories PCA of TAV and BAV lipidomes of female and male individuals across different pathophysiological stages of FCAVD. **f,** FCAVD progression is characterized by the same lipidome alterations in TAV and BAV.

Contrary to the previous observations in TAV, Lipid Trajectories PCA showed that BAV display less sex-driven dimorphism of disease progression, with both male and female trajectories situated closer to the TAV female samples in the PCA space (Fig. 4d). Again, the separation along PC2 was mainly driven by TG, CE and SP lipids in fibrotic and calcific stages of male vs female TAV (Fig. 4e).

Taken together, fibrotic and calcific TAV and BAV lipidomes share large similarities (Fig. 4f). However, BAV lipid signatures of FCAVD are not sex-specific and overall close to the pathological trajectories detected in female TAV. Of interest, however, is the specific enrichment of VLCFA SP lipids in mildly diseased BAV sections, indicating that these lipid species might be an early marker of pathological predisposition.

## Discussion

Lipid dysmetabolism is a well know driver of FCAVD onset and progression, however limited insight of the underlying mechanisms prevents further development of pharmaceutical interventions. Indeed, despite certain similarities with atherosclerosis in terms of an elevated tissue deposition of CE-rich lipoproteins, interventions based on the inhibition of the cholesterol biosynthesis pathway with statins did not show significant reduction in AS progression or severity^46^.

To support the mechanistic understanding of dyslipidemia in fibro-calcific remodeling, we performed in-depth quantitative lipidomic characterization of human AVs along FCAVD progression, covering mildly diseased, fibrotic and calcific stages. Over 1070 lipid molecular species were annotated, of which 480 were quantified. Using UMAP-based topological analysis of selected human lipidomes, we could demonstrate that the AV lipidome closely overlaps with the lipidome of blood plasma in terms of main lipoprotein-derived lipids (CE, TG, SM, PC), whereas Cer showed an AV-specific response. Lipoprotein-related molecular signatures confirmed AV enrichment with lipids from CE-rich LDL and Lp(a), complementing previously published immunohistochemistry and proteomics data^29,47^.

By mapping lipidomic signatures to the different stages of FCAVD, both extrinsic and intrinsic metabolic trajectories were identified. Lipoprotein accumulation, already evident in mildly diseased sections, peaked at the fibrotic stage and was characterized by a significant buildup of CE, free cholesterol and SP lipids, with TG having only minor impact. Importantly, the infiltration of lipoprotein-derived lipids led not just to a passive lipid deposition but was accompanied by active metabolic processing of PC to LPC and SM to Cer by the action LpPLA_2_ and SMase, respectively. Elevated LPC generally reflected the composition of the infiltrated PC precursors. On the other hand, although diverse SM species accumulated in diseased AVs, elevated Cer were mainly represented by species carrying VLCFA chains. This apparent specificity is not entirely clear and might be attributed to SP metabolism within AV cell populations in addition to the SMase action on lipoprotein-derived SM. Interestingly, the enzyme responsible for the synthesis of VLCFA Cer, ceramide synthase 2 (CerS2), is known to be highly expressed in the heart^48^.

Considering the well-established correlation between the autotaxin activity with the severity of FCAVD^12^, we looked for the endogenous levels of LPA lipids, which are often named as bioactive mediators driving FCAVD progression^49^. However, detection of endogenous LPA in AV samples processed via a conventional lipid extraction protocol was not possible. Using tailored extraction and targeted MS detection, we could quantify four LPA species of which only LPA 16:0 showed slight but significant elevation in calcific AV. Thus, we propose that LPA generation and biological activities are rather transient and hardly detectable in advanced fibrotic and calcific lesions.

Active metabolic conversion of lipoprotein-derived PC and SM to the corresponding LPC and Cer increases lipoprotein aggregation, fusion and retention in AV, representing a vicious cycle escalating lipid deposition and AV steatosis. Indeed, Cer lipids attracted much attention in cardiovascular research. Circulating levels of four particular Cer molecular species have been used as a prognostic scoring test named the Coronary Event Risk Test (CERT)^50,51^, which showed to be superior in predicting cardiovascular risk compared to traditional markers such as LDL-cholesterol (LDL-C)^52,53^. Considering the correlation between cardiovascular disease (CVD) severity and Cer-based CERT score, it is possible to speculate that Cer accumulated in AV or atherosclerotic lesions can leak back in the circulation, reflecting the valvular and vasculature status.

Interestingly, it was demonstrated that Cer composition in liver, the main site of SP synthesis and lipoprotein secreting organ, is positively correlated not only with the corresponding LDL SM species, but also with LDL aggregation^54^. Overall, the composition of hepatic TG, SP and PC lipids is closely related to that of LDL^54^. Thus, Cer synthesized in the liver are released into circulation in the form of the corresponding SM within lipoproteins, which, when trapped in AV, are converted back to the bioactive Cer species^55^. Thus, one can propose a direct link between liver dysmetabolism and FCAVD via SP lipid signatures. Indeed, correlations between AS and alcoholic and non-alcoholic fatty liver diseases was demonstrated^56–58^. Interestingly, hepatocytes-specific inhibition of CerS2 reduced levels of circulating Cer used in the CERT score^59^, further underling the significance of SP lipids in the organ-to-organ metabolic cross-talk.

In addition to FCAVD metabolic trajectories driven by lipoprotein infiltration, we illustrated AV-intrinsic lipidomic signatures. Accumulation of major membrane-building GPL at the fibrotic stage probably reflects increased cell populations due to the transformation and proliferation of AV resident cells and/or infiltrating immune cells. The calcific TAV lipidome showed degenerative signatures characterized by a massive loss of PUFA etherPE, most probably due to their oxidative degradation, and mitochondria-specific CL lipids indicating extensive cell death. Interestingly, we observed a sharp decline in the levels of PS lipids as a hallmark of progressive AV calcification. PS are one of the major anionic lipids enriched at the inner leaflet of plasma membranes and the cytosolic side of late endocytic compartments^60^. Thus, extensive PS depletion might indicate both disorganization of plasma membranes and/or cellular secretory and endocytic pathways. PS were shown as crucial players in intracellular trafficking of LDL-derived cholesterol from plasma membrane to ER^61^. In conditions of massive cholesterol overload, it is possible that PS homeostasis might be drastically dysregulated. However, the most plausible explanation is probably the role of anionic PS lipids in calcification, where negatively charged head groups of PS lipids serve as hydroxyapatite nucleator sites^62^.

Increasing evidence indicates that biological sex plays an important role in FCAVD^43–45^. Thus, one of the major findings of this study is that metabolic trajectories in FCAVD progression are highly sex-specific. Major sex-specific traits are defined by higher content of PUFA TG in male AV and elevated accumulation of SM and Cer lipids in female tissues. These sex-specific differences in the FCAVD lipidome might reflect the differences in blood lipidomes of male and female individuals^63–65^. Interestingly, in FCAVD, women are known to exhibit more fibrotic phenotypes, which can be associated with higher levels of Cer in female AVs, as identified here. SP in general and Cer in particular are emerging mediators of tissue fibrosis. The link between SP metabolism and fibrosis onset and development was shown for liver^66^, kidney^67^, cardiac^68^, and cystic fibrosis^69^. Among signaling pathways classically attributed to fibrosis development, TGF-β signaling was shown to be directly correlated with Cer levels^70^. Interestingly, TGF-β signaling is upregulated in women relative to men with AS^71,72^. Thus, based on our lipidomics data, we propose a possible mechanistic link between sex-specific TGF-β signaling, Cer accumulation and fibrosis severity, which might open a window of opportunities for therapeutic intervention to limit fibrotic AV remodeling. Interestingly, metabolic trajectories of BAV along FCAVD development were less sex-specific and generally more similar to the female TAV, thus associated with higher SP accumulation, possibly reflecting a higher prevalence of the fibrotic stage. It can also be interpreted that the hemodynamic effects are more detrimental for the pathological process in BAV, rather than the sex-specific mechanisms that may be more relevant in TAV.

In conclusion, we provide a molecular link between FCAVD development and Cer accumulation, which can be further exploited as a target of pharmacological interventions. Therapeutic strategies aiming at lowering Cer levels by inhibiting biosynthesis or enhancing their degradation present new therapeutic opportunities^36,73,74^. Thus, targeting SP rather than cholesterol metabolism might be a promising option for FCAVD treatment. Importantly, we identify significant sex-specific responses in FCAVD progression. With women presenting significantly higher levels of AV Cer, future development in SP-based diagnostics and therapeutic interventions should be performed in a sex-specific manner. Indeed, the general misperception that women are less susceptible to CVD stems from an under-representation of women in clinical trials. Even in cohorts used for the validation of CERT or sphingolipid-inclusive score (SIC), as strong CVD event predictors, 72% and 77% of the cohorts, respectively, were represented by male individuals^75,76^. Considering that a major sex-specific trait in FCAVD was related to SP metabolism, development of sex-specific diagnostic assays and treatment options must be considered.

The obvious limitation of our study is the small sample size, which would call for further validation. To this end, all lipidomics data obtained here are made publicly available to the community both as raw MS files and processed datasets, to facilitate further data interrogation and, importantly, to design targeted MS assays tailored to the human AV lipidome for high-throughput validation in larger cohorts of samples^25^.

## Methods

### Material and Methods

#### Chemicals

Acetonitrile (ULC/MS-CC/SFC grade, > 99.97%), methanol (UHPLC-MS grade, > 99.97%), isopropanol (ULC/MS-CC/SFC grade, >99.95%) and formic acid (ULC/MS-CC/SFC grade, > 99%) were obtained from Biosolve B.V. (Valkenswaald, Netherlands). Hydrochloric acid was purchased from Merck KGaA (Darmstadt, Germany). Acetic acid, ammonium formate (MS grade), butylated hydroxytoluene (BHT), chloroform (analytical grade, EMSURE®), diethyl ether, hexane, primuline and triethylamine were purchased from Sigma Aldrich Chemie GmbH (Taufkirchen, Germany). SPLASH®LIPIDOMIX®, Cer/Sph Mixture I, Cer 18:0;O2/12:0, Cer 18:0;O3/8:0, Cer 18:1;O2/17:0;O, CL 18:2/18:2/18:2/18:2 (d5), LPA 16:0, LPA 17:1, LPA 18:0, LPA 18:1, LPA 20:4, PA 17:0/17:0, PA 16:0/18:1 and TLC standards, including TG 18:0/18:0/18:0, FA 18:0, Cholesterol, DG 16:0/16:0/0:0, DG 16:0/0:0/16:0, MG 18:0/0:0/0:0, PG 16:0/18:1, PE 16:0/18:1, PI 16:0/18:1, PS 16:0/18:1, LPE 18:1, PC 16:0/18:1, SM 18:1;O2/16:0 and LPC 18:1, were obtained from Avanti Polar Lipids Inc (Alabaster, USA). Cer 18:0;O2/8:0 was from Sigma Aldrich. Water (resistance R > 18 MΩ/cm, total organic content less than 10 ppb) was purified on a PureLab Ultra Analytic system (ELGA Lab water, Celle, Germany) or obtained from Biosolve B.V. (ULC/MS-CC/SFC grade, Valkenswaald, Netherlands). Optimum cutting temperature compound was purchased from Sakura Finetek USA (Torrance, CA, USA). Movat Pentachrom Kit was purchased from Morphisto GmbH (Offenbach am Main, Germany). ROTI Histokitt II was purchased from Carl Roth GmbH + Co. KG (Karlsruhe, Germany).

### Human aortic valve tissue samples

FCAVD tissues were obtained from 55 patients undergoing aortic valve replacement for severe AS (Supplementary Data Table 1). Tissue collection was approved by the local Ethical Committee (Medical Faculty, University Leipzig, registration number 128/19-ek) and all patients gave written informed consent in accordance with the Declaration of Helsinki. All samples showed macroscopic calcification as a sign of progressed FCAVD. Tissue samples were dissected into mildly diseased, fibrotic and calcific sections on-site and flash frozen in liquid nitrogen (Fig. 2a). Deep-frozen tissue sections grinded in a percussion mortar in liquid nitrogen and stored at -80 °C until further analysis.

### Pentachrome staining

Native AV samples were embedded perpendicularly to obtain longitudinal sections in OCT Embedding Matrix for Frozen Sections (Cell Path, Carl Roth #6478.1). Consecutive sections of 5 µm were cut using Cryostat (Leica CM1860, Germany) and mounted onto microscope slides (Superfrost Plus Slides, Epredia, USA). Cryosections were dried and fixed in Methanol/Aceton 1:1 solution. Movat Pentachrom Kit (Morphisto, Germany) was used according to the manufacturer’s instructions. Following this staining nuclei appear in blue-black, cytoplasm in light red, elastic and muscle fibers in red, collageneous connective tissue in light yellow, mineralized bone in dark yellow and mineralized cartilage tissue in blue-green. The sections were mounted with xylene-containing mounting medium (ROTI Histokit II from Carl Roth #T160.1).

To quantify calcification status of the samples sections were incubated in phosphate buffered saline containing 100 pmol/ml Osteosense solution (IVI SenseOsteo 680 #NEV10020Ex, PerkinElmer) for 24h at 37 °C in the dark. Then, Hoechst 33342 (Miltenyi, Bergisch-Gladbach, Germany) was added in a 1000-fold dilution and incubated for another ten minutes. Finally, sections were washed with water and the slides were mounted using Roti Mount FluorCare (Carl Roth #HP19.1). Pictures were taken using Keyence BZ-X800E (Keyence, Germany).

### Lipid extraction of pooled AV sample for lipid identification and ITSD design

All solvents used for the extraction were supplemented with 0.1 % (w/v) BHT and cooled on ice before use. All extraction steps were performed on ice or at 4 °C. Pooled sample was formed by mixing equal amounts of AV tissue powders from each specimen (mildly diseased, fibrotic and calcific). Lipids were extracted according to the Folch protocol^77^. 20 mg of pooled sample was placed into 2 mL tubes (Eppendorf, Hamburg, Germany) and 600 µL chloroform/methanol (2:1, v/v) was added. Samples were incubated for 1 h at 4 °C on a rotary shaker (300 rpm). 150 µL water was added to induce phase separation and mixture was incubated for another 15 min at 4 °C on a rotary shaker (300 rpm). Samples were centrifuged (10,000 × g, 10 min, 4 °C), the lower organic phase was transferred to a new tube and dried under vacuum.

### Untargeted lipidomics workflow for human AV lipids identification

Lipid extract obtained from the pooled AV sample was reconstituted in 200 µL isopropanol. 5 µL (for analysis in positive ionization mode) and 10 µL (for analysis in negative ionization mode) were loaded on an Accucore reverse phase C30 column (2.1 × 150 mm, 2.6 μm, 150 Å; Thermo Fisher Scientific) installed on a Vanquish Horizon UHPLC (Thermo Fisher Scientific). UHPLC separation was coupled online either to Q Exactive Plus Hybrid Quadrupole Orbitrap (ID Method 1, 34 min gradient) or Exploris 240 Hybrid Quadrupole Orbitrap (ID Method 2, 57 min gradient) mass spectrometers both equipped with a HESI source (Thermo Fisher Scientific).

#### ID Method 1, 34 min gradient

Lipids were separated by gradient elution with solvent A (acetonitrile/water, 1:1, v/v) and B (isopropanol/acetonitrile/water, 85:10:5, v/v/v) both containing 5 mM ammonium formate and 0.1% (v/v) formic acid. Separation was performed at 50 °C with a flow rate of 0.3 mL/min using the following gradient: 0-20 min – 10 to 86% B, 20-22 min – 86 to 95% B, 22-26 min – 95% B, 26-26.1 min – 95 to 10% B, followed by 7.9 min re-equilibration at 10% B. Mass spectra were acquired in positive and negative ionization modes using spray voltages of 3.5 kV and - 2.5 kV, respectively, ion transfer temperature of 300 °C, aux gas heater temperature of 370 °C, S-lens RF level of 35%, auxiliary gas of 10 a.u., sheath gas of 40 a.u., and sweep gas of 1 a.u.. In positive ion mode, full scan mass spectra were recorded from *m*/*z* 250 to 1200 at a resolution of 140 000 at *m*/*z* 200, AGC target of 1×10^6^, maximum injection time of 100 ms, and default charge state of 1. In negative ion mode scan ranges from *m*/*z* 380 to 1200 or from *m/z* 400 to 1600 were applied. Tandem mass spectra were recorded by data-dependent acquisition (DDA) for the 15 most intense precursor ions (DDA top 15) at a resolution of 17 500 at *m*/*z* 200, AGC target of 1×10^5^, maximum injection time of 60 ms (negative mode: 150 ms), isolation window of 1.2 *m*/*z* units, and stepped normalized collision energy (NCE; 10, 20, and 30). The DDA settings were set to an apex trigger of 6 s, isotope exclusion on, and dynamic exclusion for 10 s^26^.

#### ID Method 2, 57 min gradient

Lipids were separated by gradient elution with solvent A (acetonitrile/water, 1:1, v/v) and B (isopropanol/acetonitrile/water, 85:10:5, v/v/v) both containing 5 mM ammonium formate and 0.1% (v/v) formic acid. Separation was performed at 50 °C with a flow rate of 0.3 mL/min using following gradient: 0-20 min – 10 to 80% B, 20-37 min – 80 to 95% B, 37-41 min – 95% to 100% B, 41-49 min – 100% B, 49-49.1 100% to 10% B, followed by 7.9 min re-equilibration at 10% B. Mass spectra were acquired in positive and negative ionization modes using spray voltages of 3.5 kV and - 2.5 kV, respectively, ion transfer temperature of 300 °C, aux gas heater temperature of 370 ◦C, S-lens RF level of 35%, auxiliary gas of 10 a.u., sheath gas of 40 a.u., and sweep gas of 1 a.u. In positive ion mode, full scan mass spectra were recorded from *m*/*z* 200 to 1200 at a resolution of 120 000 at *m*/*z* 200, AGC target of 1×10^6^, maximum injection time was set to auto. In the negative ion mode, scan ranges from *m/z* 200 to 1400 and *m*/*z* 370 to 1480 with a maximum injection time of 100 ms were acquired. Easy-IC was set to RunStart for all methods. Tandem mass spectra were recorded by data-dependent acquisition (DDA) within a cycle time of 1.3 s at a resolution of 30000 at *m*/*z* 200, an AGC target of 1×10^5^, a maximum injection time of 54 ms, an isolation window of 1.2 *m*/*z* units, a stepped NCE (17, 27, 37), and charge state 1. The DDA settings were set to isotope exclusion on, dynamic exclusion for 6 s (mass tolerance ±2.5 ppm, exclusion after 2 times within 10 s).

Software-assisted lipid identification was performed using Lipid Hunter^78^ (source code version: https://github.com/SysMedOs/lipidhunter). RAW files were converted with MSconvert (ProteoWizard version 3.0.9134) into mzML format. Lipids were identified using a mass accuracy of 5 and 20 ppm for MS1 and MS/MS spectra, respectively. The remaining parameters were kept as default. Proposed lipid species were confirmed by manual inspection of the tandem mass spectra for species-specific fragment ions within the HTML report. CE, CL and GM3 gangliosides were identified manually. Proposed identifications were further validated by plotting the retention time of lipid species against their Kendrick mass defect by hydrogen and lipid species not following expected trendlines were excluded.

### Design of the AV-tailored mixture of internal standards (ISTD) for semi-absolute quantification

To design ISTD mixture tailored to the lipid subclasses and their endogenous concentrations in human AV lipidome, the rough composition and relative abundance of AV lipids was first accessed via quantitative high performance thin layer chromatography (qHPTLC). To this end, pooled AV lipid extract (prepared as described above) was dissolved in 200 µL chloroform/methanol (2:1, v/v) and loaded onto a TLC plate (10 or 20 µL, HPTLC silica gel 60, 20 × 10 cm, Merck) using a Camag Linomat 5 sampler (Camag, Switzerland). On each plate six-point serial dilutions of polar or apolar lipid TLC standards were loaded for quantitative lipid class-specific calibration (Supplementary Data Table 4). Plates were developed using chloroform/ethanol/trimethylamine/water (5:5:5:1, v/v/v/v) or hexane/diethylether/acetic acid (85:15:1, v/v/v/v) for polar and apolar lipid quantification, respectively. Dried TLC plates were immersed in acetone/water (8:2, v/v) containing primuline (0.05 %, w/v) for 5 s (Camag Chromatogramm Immersion Device III). Analytes were visualized under UV light (366 nm), and scanned by a videodensitometric device (Biostep GmbH, Germany). Densitometric analysis was performed with Image Lab (Version 6.1, Bio-Rad). Using TLC-based quantification (Supplementary Data Figs. 1a and b), the rough abundances of major AV lipid classes were approximated, based on which the initial composition of ISTD mixture was designed by combining SPLASH® Lipidomix® : Cer/SpH Mix I in the ratio 1:1,5 (v/v), and additional CL and individual Cer standards were added (Supplementary Data Table 5).

For defining the appropriate amounts of ISTD mixture to be spiked into individual samples for semi-absolute quantification via single-point calibration, seven-point calibration curves were generated. To this end, pooled AV tissue powder was spiked (each calibration point in independent triplicates) with selected ISTD mixture in seven different amounts (Supplementary Data Table 5). 21 samples were extracted as described above, extracts were reconstituted in 100 µL isopropanol, and analysed by LC-MS using C30 RPLC separation coupled online to Q Exactive Plus Hybrid Quadrupole Orbitrap mass spectrometer as described above with slight adaptations. Individual samples were recorded in full scan (MS1) positive and negative ion modes in the range from *m*/*z* 100 to 1500. Additionally, 10 µL of each sample was combined into a total quality control (tQC) sample, which was acquired in DDA as described above in order to map lipid annotations to full scan mode (MS1) data when necessary. Peak areas of five to nine endogenous lipids per subclass (selected from the identification Table S2) representing the highest, the middle and the least abundant species as well as the corresponding ISTD were integrated using Skyline (version 22.2) software^79^. Area under curve (AUC) for different class-specific adducts and common in-source fragments (see below for more details) were summed up. Incomplete isotopic enrichment was corrected for deuterated ISTD. All peaks were subjected to type I isotopic corrections^80^.

For each ISTD seven points calibration curves were generated (Supplementary Data Fig. 2) and linearity range of LC-MS responses was accessed. Additionally, peak areas for each ISTD were plotted relative to the peak areas of the endogenous AV lipids to select the ISTD amounts not only within the linear range of LC-MS responses but also preferentially representing the middle range of the endogenous lipids concentrations for each considered lipid subclass (Supplementary Data Fig. 3).

Final AV-tailored ISTD mixture was prepared by combining individual standards from the stock solutions in the required amounts (Supplementary Data Table 6). Complete ISTD mixture was prepared once in the quantities sufficient for all individual samples, dried under vacuum, re-dissolved in chloroform/methanol (2:1, v/v) and divided into working aliquots (one per extraction batch) that were dried, stored at -80 °C, and reconstituted in chloroform/methanol (2:1, v/v) before use. To each sample corresponding roughly to 20 mg of AV tissue, 10 µL of ISTD mixture with individual standards amount provided in Supplementary Data Table 6 was added before the lipid extraction.

### Extraction and untargeted lipidomics workflow for human AV lipids quantification

All individual samples were extracted and analysed in a randomized order to account for possible batch effects during sample preparation and LC-MS experiments. For the evaluation of potential batch effects, representative pools were created by combining sample material from the same sample cohort (batch quality control, BQC). Samples were randomly grouped into batches of a maximum of 35 samples per batch. The BQC samples were split into aliquots of 20 mg each and distributed to the different batches. Samples and BQCs of the same batch were extracted simultaneously. All solvents used for extraction were supplemented with 0.1 % (w/v) BHT and cooled on ice before use. All extraction steps were performed on ice or at 4 °C. Samples were allowed to thaw on ice for 30 min prior to extraction. To approximately 20 mg of each individual AV sample tissue powders 600 µL chloroform/methanol (2:1, v/v) and 10 μL of ISTD mixture in chloroform/methanol (2:1, v/v; Supplementary Data Table 6) were added, mixed and incubated on ice for 15 min. Samples were incubated for 1 h at 4 °C on a rotary shaker (300 rpm). 150 µL water was added to induce phase separation and the mixture was incubated for 15 min at 4 °C on a rotary shaker (300 rpm). Samples were centrifuged (10,000 × g, 10 min, 4 °C), the lower organic phase was transferred to a new tube, and dried under vacuum.

Lipid extracts were reconstituted in 110 µL isopropanol and 2 µL were loaded on an Accucore reverse phase C30 column (2.1 × 150 mm, 2.6 μm, 150 Å; Thermo Fisher Scientific) installed on a Vanquish Horizon UHPLC (Thermo Fisher Scientific) coupled online to an Exploris 240 Hybrid Quadrupole Orbitrap mass spectrometer. Lipids were separated by gradient elution with solvent A (acetonitrile/water, 1:1, v/v) and B (isopropanol/acetonitrile/water, 85:10:5, v/v/v) both containing 5 mM ammonium formate and 0.1% (v/v) formic acid. Separation was performed at 50 °C with a flow rate of 0.3 mL/min using following gradient: 0-10 min – 30 to 80% B, 10-27 min – 80 to 95% B, 27-31 min – 95% to 100% B, 31-39 min – 100% B, 39-39.1 100% to 10% B, followed by 7.9 min re-equilibration at 10% B. Mass spectra were acquired in positive mode using spray voltages of 3.5 kV, ion transfer temperature of 300 °C, aux gas heater temperature of 370 °C, S-lens RF level of 35%, auxiliary gas of 10 a.u., sheath gas of 40 a.u., and sweep gas of 1 a.u. Easy-IC was set to RunStart. Mass spectra were recorded in full scan (MS1) from *m*/*z* 200 to 1200 at the resolution of 120 000 at *m*/*z* 200, AGC target of 1×10^6^, maximum injection time auto. Additionally, 10 µL of each sample was combined into a total quality control (tQC) sample, which was acquired after every 10 samples as well as at the start of the sequence in DDA mode (positive and negative ionization modes; described above) in order to map lipid annotations to full scan mode (MS1) data if necessary.

Lipids were quantified using Skyline (version 22.2)^79^. Area under curve (AUC) for different class-specific adducts and common in-source fragments (Supplementary Data Table 7) were summed up. Incomplete isotopic enrichment was corrected for deuterated ISTD. All signals were subjected to type I isotopic corrections^80^. Individual lipid species were quantified in relation to the respective lipid class specific ISTD and normalized to the wet tissue weight. Linear regression analysis was applied by plotting the calculated concentrations against their AUC value to identify and exclude possible outliers and features showing non-linear behaviour.

### Assessment of PA and LPA recovery rates

To determine the recovery of PA and LPA lipids, pooled AV tissue powders were spiked with selected PA/LPA ISTD (LPA 18:1, PA 17:0/17:0, PA 18:0/18:0, PA 18:0/18:2) with the amounts corresponding to 0, 10, 100, and 1000 pmol per 20 mg of tissue (in triplicates) before and post lipid extraction. Lipid extraction was performed using a chloroform/methanol/water mixture (Folch extraction; as described above) or by acidified methanol/chloroform extraction^38^. Briefly, for acidified methanol/chloroform extraction, 20 mg of pooled AV were mixed with 800 µL ice cold methanol/0.1 M HCl in water (1:1; v/v). 10 μL of PA/LPA ISTD in chloroform/methanol (2:1, v/v) were added, mixed and incubated for 15 min on ice. Next, phase separation was induced by adding 400 µL of chloroform, samples were centrifuged (10,000 × g, 10 min, 4 ◦C), lower organic phase transferred to a new tube and dried under vacuum. All solvents used for the extraction were supplemented with 0.1 % (w/v) BHT and cooled on ice before use. All extraction steps were performed on ice.

Lipid extracts were reconstituted in 100 µL methanol/isopropanol (1/1, v/v), 5 µL were loaded on an Accucore reverse phase C30 column (2.1 × 150 mm, 2.6 μm, 150 Å; Thermo Fisher Scientific) installed on a Vanquish Horizon UHPLC (Thermo Fisher Scientific). Lipids were separated by gradient elution with solvent A (acetonitrile/water, 1:1, v/v) and B (isopropanol/acetonitrile/water, 85:10:5, v/v/v) both containing 5 mM ammonium formate and 0.1% (v/v) formic acid. Separation was performed at 50 °C with a flow rate of 0.3 mL/min using following gradient: 0-20 min – 10 to 86% B, 20-22 min – 86 to 95% B, 22-26 min – 95% B, 26-26.1 min – 95% to 10% B, followed by 7.9 min re-equilibration at 10% B. Mass spectra were acquired on Exploris 240 operating in negative ionization modes using spray voltage of - 2.5 kV, ion transfer temperature of 300 °C, aux gas heater temperature of 370 °C, S-lens RF level of 35%, auxiliary gas of 10 a.u., sheath gas of 40 a.u., and sweep gas of 1 a.u. Easy-IC was set to RunStart. Full scan mass spectra were recorded from *m*/*z* 200 to 1200 at a resolution of 120 000 at *m*/*z* 200, AGC target of 1×10^6^, maximum injection time of 100 ms, and default charge state of 1. Tandem mass spectra were recorded by data-dependent acquisition (DDA) within a cycle time of 1.3 s at a resolution of 15 000 at *m*/*z* 200, an AGC target of 1×10^5^, a maximum injection time of 60 ms, an isolation window of 1.2 *m*/*z* units, and a stepped NCE (17, 27, 37), isotope exclusion on, and dynamic exclusion for 6 s. Lipids were quantified using Skyline (version 22.2) as described above. Recovery rates were determined by dividing the mean AUC of the samples spiked with PA/LPA ISTD before and post extraction multiplied by 100%.

### Extraction and targeted lipidomics workflow for human AV lipids quantification

For quantification of LPA, LPI, LPS, PA, PI, PS, PG, CL, and GM3 lipids acidified extraction and targeted LC-MS/MS analysis (selected reaction monitoring, SRM; see details below) in negative ionization mode was used. First, to determine the appropriate amounts of ISTD, a seven-point calibration curves were generated using workflow described above (Supplementary Data Table 5). Linearity of the response was evaluated and final amount of ISTD to be spiked in 20 mg of AV tissue powder were selected (Supplementary Data Table 6). Approximately 20 mg of each individual sample (Supplementary Data Table 1) was spiked with selected ISTD amounts and used for the acidified extraction as described above. Lipid extracts were reconstituted in 100 µL methanol/isopropanol (1:1, v/v) and 3 µL were loaded on an Accucore reverse phase C18 column (2.1 × 150 mm, 2.6 μm, 80 Å; Thermo Fisher Scientific) installed on a Vanquish Flex UHPLC (Thermo Fisher Scientific) coupled online to TSQ Altis Plus Triple Quadrupole mass spectrometer equipped with a HESI source (Thermo Fisher Scientific). Lipids were separated by gradient elution with solvent A (acetonitrile/water, 1:1, v/v) and B (isopropanol/acetonitrile/water, 85:10:5, v/v/v) both containing 5 mM ammonium formate and 0.1% (v/v) formic acid. Separation was performed at 50 °C with a flow rate of 0.3 mL/min using following gradient: 0-7 min – 30 to 85% B, 7-8 min – 85 to 95% B, 8-10 min – 95% B, 10-10.1 min – 95% to 30% B, followed by 4.9 min re-equilibration at 10% B. Mass spectra were acquired in selected reaction monitoring (SRM) mode using spray voltages of -2.5 kV, ion transfer temperature of 300 ◦C, vaporizer temperature of 370 ◦C, auxiliary gas of 10 a.u., sheath gas of 50 a.u., and sweep gas of 1 a.u.. SRMs were recorded setting automatic dwell time to ensure acquisition of 10 points per peak, dwell time factor 3, Q1 and Q3 resolution (FWHM) of 0.7 Da, and CID gas flow of 1.5 mTorr. Lipid-species specific transitions, collision energy, and RF lens voltage for each analyte are listed in the Supplementary Data Table 8. For quantification, data were processed using Skyline (version 22.2) as described above. Both type I and II isotopic corrections were applied and individual lipid species were quantified by the respective lipid class specific ISTD and normalized to the sample weight. Linear regression analysis was applied by plotting the calculated concentration of lipid species against their AUC value to identify and exclude outliers and features with non-linear behaviour.

### Statistical analysis

Quantitative data analysis including isotopic corrections, ISTD normalization and normalization by sample weight was performed with Microsoft Excel 2016. Graphical representations were generated using Corel DRAW 2020 22.0 (Corel corporation), Graphpad Prism® 8.0.2 (GraphPad Software, Inc.), Inkscape 1.3.2 (Inkscape Developers), Python 3.12 (Python Software Foundation) and R Statistical Software v4.4.1(R Core Team 2021). Statistical analysis was performed using Graphpad Prism® 8.0.2, MetaboAnalyst version 6.0^81^, Python 3.12 and R Statistical Software v4.4.1. Statistical significance between compared groups were found by t-test or with an ANOVA with a threshold of p ≤ 0.05. Generic Python libraries for data processing and visualization tasks such as Pandas, Scikit-learn, Matplotlib, and Plotly are used in different plots. The detailed configurations and versions can be found in the corresponding source code repositories.

### Lipid distribution plot

The lipid quantification results were averaged and used for the generation of lipid distribution plot using customized Python scripts. Inspired by the distribution plot published by Bo Burla et.al^82^, a python version of the distribution plot was developed and used for the successful visualization of adipocyte’s lipidome for the AdipoAtlas project^26^ (https://github.com/SysMedOs/AdipoAtlasScripts). The code takes the concentration of each lipid, plot a colour bar on the log scale according to corresponding lipid subclass. A bold colour bar representing the total sum of each lipid subclass is plotted automatically. Based on the previous code base, the improved version to visualize the distribution of lipids is summarized into the LipidomeDistribution repository and the source code is released under AGPL v3 license on GitHub: https://github.com/LMAI-TUD/LipidomeDistribution.

### Uniform manifold approximation and projection (UMAP) algorithm

To generate UMAP, lipid quantities, fatty acyl chains composition (carbon and double bound count) and myopic MCES distances for lipid class specific structures were considered. Briefly, SMILES representation of the lipid class specific backbone (lipid_class_backbone_smiles) without FA residues were generated and the myopic MCES distances to other lipid_class_backbone_smiles were calculated using myopic-mces package^83^ (source code: https://github.com/boecker-lab/myopic-mces). Resulted myopic MCES distances matrix (24×24) was used to perform a PCA (number of components = 10) to reduce data dimensionality and the top 5 components were further used for UMAP analysis. To bring all used values in the similar range, lipid concentrations were z-scored, and all other values were scaled between 0 and 1 using min-max scaler by Scikit-learn Python library. Different weight factors were applied to the scaled values to generate a final data matrix for the UMAP representation. A detailed description of all parameters was summarized in the Supplementary Data Table 9. The source code of LipidomeUMAP is released under AGPL v3 license on GitHub: https://github.com/LMAI-TUD/LipidomeUMAP.

### Sankey plot

Sankey plot for the visualization of Sphingolipids was previously introduced by the AdipoAtlas project^26^ (https://github.com/SysMedOs/AdipoAtlasScripts). In brief, for each sphingolipid, the sphingoid base, the FA residue, and the corresponding sphingolipids sub class is assigned and connected in the concept of the Sankey flow diagram. The widely used visualization library Plotly for Python was used to generate the Sankey plot from the identified sphingolipids. The updated Sankey plot for this project is released under AGPL v3 license on GitHub: https://github.com/LMAI-TUD/LipidSankey.

### Lipid Trends Analysis

Trend analysis diagrams was generated by plotting the individual lipid clusters from Gaussian Mixture Model clustering of lipids according to their concentration variances accross the mildly diseased, fibrotic and calcific stages. The missing values in the quantified data matrix table were filled with 1/5 of the minimum value of the corresponding lipid across all samples according to the widely used MetaboAnalyst tool^81^. The filled data matrix was averaged to the mildly diseased, fibrotic, calcific stages. The averaged data matrix was performed the Z-score scaling followed by Gaussian Mixture Model clustering using the mainstream scikit-learn Python library. The clustered results were then plotted as trend plot for each cluster by matplotlib Python library. After evaluation of different combination of scikit-learn built-in scaling methods (min-max, Z-score, and Log2) with mainstream clustering algorithms (Hierarchical clustering, K-means, bisecting K-means, Gaussian Mixture Model, and Dirichlet Process Gaussian Mixture Model), the Z-score and Gaussian Mixture Model combination was selected for the generation of the plots. The reusable python scripts providing generic access to multiple scaling and clustering algorithms are provided on GitHub under AGPL v3 license: https://github.com/LMAI-TUD/LipidTrends.

### Trajectories PCA plot

The quantified data matrix with metadata labels for the sample ID, groups (TAV male, TAV female), and sections (mildly diseased, fibrotic, calcific) was used for the generation of the trajectory PCA plot. The trajectory PCA visualization is inspired by the clinical lipidomics analysis R script from Singapore Lipidomics Incubator (SLING) (https://github.com/SLINGhub/iSLS12_2024). The missing values were filled with 1/5 of the minimum value of the corresponding lipid across all samples according to the widely used MetaboAnalyst tool^81^. The filled data matrix was averaged to the defined groups and progression stages. After log2 scaling, the averaged and scaled data matrix was then used for PCA with a number of components set to 5. The trajectory arrows on the PCA scores plots visually highlight the trends from the mildly diseased to fibrotic and calcific stage on the 2D space of PC1 and PC2 of the PCA results. The scripts used for the plot can be easily modified to be compatible with other datasets and adjust the main parameters such as the missing value filling method, scaling methods (Z-sore or log2) and number of the PCA components. The TrajectoryPCA source code is released under AGPL v3 license on GitHub: https://github.com/LMAI-TUD/TrajectoryPCA.

## Data availability

The data used for above mentioned plots can be accessed in corresponding GitHub code repositories from the LMAI team https://github.com/LMAI-TUD. The raw data/mzML spectra and processed lipidomics data can be accessed through Metabolomics Workbench ST003364 (doi: 10.21228/M8G24F)25.

## Code availability

The source code used for the above mentioned plots can be accessed in the corresponding GitHub code repositories from the LMAI team https://github.com/LMAI-TUD.

## Author contributions

Conceptualization and study design was carried out by P.P., J.S., F.S. and M.F.. Initial collection of sample material and preparation was performed by J.B., S.W., H.T., P.B. and F.S.. Sample preparation for mass spectrometry experiments and data acquisition was performed by P.P. and M.W.. J.B., S.W. and P.B. performed histological staining. Analysis and interpretation of data was carried out by P.P., M.W., Z.N., and M.F.. P.P. and M.F. wrote the manuscript with contributions from all co-authors.

## References

1. Goody, P. R. et al. Aortic valve stenosis: from basic mechanisms to novel therapeutic targets. Arterioskler. Throm. Vasc. 40, 885–900 (2020); 10.1161/ATVBAHA.119.313067.

2. Everett, R. J. et al. Progression of hypertrophy and myocardial fibrosis in aortic stenosis: a multicenter cardiac magnetic resonance study. JACC Cardiovasc. Imaging 11, e007451 (2018); 10.1161/CIRCIMAGING.117.007451.

3. Nkomo, V. T. et al. Burden of valvular heart diseases: a population-based study. Lancet 368, 1005–1011 (2006); 10.1016/S0140-6736(06)69208-8.

4. Small, A. M., et al. Unraveling the mechanisms of valvular heart disease to identify medical therapy targets: a scientific statement from the American heart association. Circulation; 10.1161 (2024), in press. CIR.0000000000001254.

5. Matilla, L. et al. Sex-differences in aortic stenosis: mechanistic insights and clinical implications. Front. Cardiovas. Med. 9, 818371 (2022); 10.3389/fcvm.2022.818371.

6. Otto, C. M., Kuusisto, J., Reichenbach, D. D., Gown, A. M. & O’Brien, K. D. Characterization of the early lesion of ‘degenerative’ valvular aortic stenosis. Histological and immunohistochemical studies. Circulation 90, 844– 853 (1994); 10.1161/01.cir.90.2.844.

7. Pohle, K. et al. Progression of aortic valve calcification: association with coronary atherosclerosis and cardiovascular risk factors. Circulation 104, 1927–1932 (2001); 10.1161/hc4101.097527

8. Smith, J. G. et al. Association of low-density lipoprotein cholesterol-related genetic variants with aortic valve calcium and incident aortic stenosis. JAMA 312, 1764–1771 (2014); 10.1001/jama.2014.13959.

9. Kaltoft, M., Langsted, A. & Nordestgaard, B. G. Triglycerides and remnant cholesterol associated with risk of aortic valve stenosis: Mendelian randomization in the Copenhagen General Population Study. Eur. Heart J. 41, 2288–2299 (2020); 10.1093/eurheartj/ehaa172.

10. Nazarzadeh, M. et al. Plasma lipids and risk of aortic valve stenosis: a Mendelian randomization study. Eur. Heart J. 41, 3913–3920 (2020); 10.1093/eurheartj/ehaa070.

11. Umezu-Goto, M. et al. Autotaxin has lysophospholipase D activity leading to tumor cell growth and motility by lysophosphatidic acid production. Cell Biol. 158, 227–233 (2002); 10.1083/jcb.200204026.

12. Bouchareb, R. et al. Autotaxin derived from lipoprotein(a) and valve interstitial cells promotes inflammation and mineralization of the aortic valve. Circulation 132, 677–690 (2015); 10.1161/CIRCULATIONAHA.115.016757.

13. Mahmut, A. et al. Elevated expression of lipoprotein-associated phospholipase A2 in calcific aortic valve disease: implications for valve mineralization. J. Am. Coll. Cardiol. 63, 460–469 (2014); 10.1016/j.jacc.2013.05.105.

14. Schissel, S. L. et al. Secretory sphingomyelinase, a product of the acid sphingomyelinase gene, can hydrolyze atherogenic lipoproteins at neutral pH. Implications for atherosclerotic lesion development. J. Biol. Chem. 273, 2738–2746 (1998); 10.1074/jbc.273.5.2738.

15. Oörni, K., Hakala, J. K., Annila, A., Ala-Korpela, M. & Kovanen, P. T. Sphingomyelinase induces aggregation and fusion, but phospholipase A2 only aggregation, of low density lipoprotein (LDL) particles. Two distinct mechanisms leading to increased binding strength of LDL to human aortic proteoglycans. J. Biol. Chem. 273, 29127–29134 (1998); 10.1074/jbc.273.44.29127.

16. Côté, C. et al. Association between circulating oxidised low-density lipoprotein and fibrocalcific remodelling of the aortic valve in aortic stenosis. Heart 94, 1175–1180 (2008); 10.1136/hrt.2007.125740.

17. Li, F., Zhao, Z., Cai, Z., Dong, N. & Liu, Y. Oxidized low-density lipoprotein promotes osteoblastic differentiation of valvular interstitial cells through RAGE/MAPK. Cardiology 130, 55–61 (2015); 10.1159/000369126.

18. Rossebø, A. B. et al. Intensive lipid lowering with simvastatin and ezetimibe in aortic stenosis. *New Engl*. J. Med. 359, 1343–1356 (2008); 10.1056/NEJMoa0804602.

19. Cowell, S. J. et al. A randomized trial of intensive lipid-lowering therapy in calcific aortic stenosis. *New Engl*. J. Med. 352, 2389–2397 (2005); 10.1056/NEJMoa043876.

20. Chan, K. L., Teo, K., Dumesnil, J. G., Ni, A. & Tam, J. Effect of Lipid lowering with rosuvastatin on progression of aortic stenosis: results of the aortic stenosis progression observation: measuring effects of rosuvastatin (ASTRONOMER) trial. Circulation 121, 306–314 (2010); 10.1161/CIRCULATIONAHA.109.900027.

21. Surendran, A. et al. Metabolomic signature of human aortic valve stenosis. JACC Basic Transl. Sci. 5, 1163– 1177 (2020); 10.1016/j.jacbts.2020.10.001.

22. Lim, J. et al. Lipid mass spectrometry imaging and proteomic analysis of severe aortic stenosis. J. Mol. Hist. 51, 559–571 (2020); 10.1007/s10735-020-09905-5.

23. Lehti, S. et al. Modified lipoprotein-derived lipid particles accumulate in human stenotic aortic valves. PloS One 8, e65810 (2013); 10.1371/journal.pone.0065810.

24. Sud, M. et al. Metabolomics Workbench: An international repository for metabolomics data and metadata, metabolite standards, protocols, tutorials and training, and analysis tools. Nucl. Acids Res. 44, D463–70 (2016); 10.1093/nar/gkv1042.

25. Prabutzki, P. et al. Deep lipidomic profiling reveals sex dimorphism of lipid metabolism in fibro-calcific aortic valve disease data sets. Metabolomics Workbench 10.21228/M8G24F (2024).

26. Lange, M. et al. AdipoAtlas: A reference lipidome for human white adipose tissue. Cell Rep. Med. 2, 100407 (2021); 10.1016/j.xcrm.2021.100407.

27. Vvedenskaya, O. et al. Nonalcoholic fatty liver disease stratification by liver lipidomics. J. Lipid Res. 62, 100104 (2021); 10.1016/j.jlr.2021.100104.

28. Schlotter, F. et al. ApoC-III is a novel inducer of calcification in human aortic valves. J. Biol. Chem. 296, 100193 (2021); 10.1074/jbc.RA120.015700.

29. O’Brien, K. D. et al. Apolipoproteins B, (a), and E accumulate in the morphologically early lesion of ‘degenerative’ valvular aortic stenosis. Arterioscler. Thromb. Vasc. Biol. 16, 523–532 (1996); 10.1161/01.atv.16.4.523.

30. Tsuji, T., Yuri, T., Terada, T. & Morita, S.-Y. Application of enzymatic fluorometric assays to quantify phosphatidylcholine, phosphatidylethanolamine and sphingomyelin in human plasma lipoproteins. Chem. Phys. Lipids 238, 105102 (2021); 10.1016/j.chemphyslip.2021.105102.

31. Ruuth, M. et al. Susceptibility of low-density lipoprotein particles to aggregate depends on particle lipidome, is modifiable, and associates with future cardiovascular deaths. Eur. Heart J. 39, 2562–2573 (2018); 10.1093/eurheartj/ehy319.

32. Skipski, V. P. et al. Lipid composition of human serum lipoproteins. Biochem. J. 104, 340–352 (1967); 10.1042/bj1040340.

33. Wiesner, P., Leidl, K., Boettcher, A., Schmitz, G. & Liebisch, G. Lipid profiling of FPLC-separated lipoprotein fractions by electrospray ionization tandem mass spectrometry. J. Lipid Res. 50, 574–585 (2009); 10.1194/jlr.D800028-JLR200.

34. Dzobo, K. E. et al. Diacylglycerols and lysophosphatidic acid, enriched on lipoprotein(a), contribute to monocyte inflammation. Arterioscler. Thromb. Vasc. Biol. 44, 720–740 (2024); 10.1161/ATVBAHA.123.319937.

35. Holčapek, M. et al. Determination of nonpolar and polar lipid classes in human plasma, erythrocytes and plasma lipoprotein fractions using ultrahigh-performance liquid chromatography-mass spectrometry. J. Chromatogr. A 1377, 85–91 (2015); 10.1016/j.chroma.2014.12.023.

36. Zietzer, A., Düsing, P., Reese, L., Nickenig, G. & Jansen, F. Ceramide Metabolism in Cardiovascular Disease: A Network With High Therapeutic Potential. Arterioscler. Thromb. Vasc. Biol.42, 1220–1228 (2022); 10.1161/ATVBAHA.122.318048.

37. Torzewski, M. et al. Lipoprotein(a) associated molecules are prominent components in plasma and valve leaflets in calcific aortic valve stenosis. JACC Basic Transl. Sci. 2, 229–240 (2017); 10.1016/j.jacbts.2017.02.004.

38. Triebl, A. et al. Quantitation of phosphatidic acid and lysophosphatidic acid molecular species using hydrophilic interaction liquid chromatography coupled to electrospray ionization high resolution mass spectrometry. J. Chromatogr. A 1347, 104–110 (2014); 10.1016/j.chroma.2014.04.070.

39. Zhao, Z. & Xu, Y. Measurement of endogenous lysophosphatidic acid by ESI-MS/MS in plasma samples requires pre-separation of lysophosphatidylcholine. J. Chromatogr. B Analyt. Technol. Biomed. Life Sci. 877, 3739–3742 (2009); 10.1016/j.jchromb.2009.08.032.

40. Sens, A. et al. Pre-analytical sample handling standardization for reliable measurement of metabolites and lipids in LC-MS-based clinical research. J. Mass Spectrom. Adv. Clin. Lab. 28, 35–46 (2023); 10.1016/j.jmsacl.2023.02.002.

41. Onorato, J. M. et al. Challenges in accurate quantitation of lysophosphatidic acids in human biofluids. J. Lipid Res. 55, 1784–1796 (2014); 10.1194/jlr.D050070.

42. Glaros, E. N., Kim, W. S., Rye, K.-A., Shayman, J. A. & Garner, B. Reduction of plasma glycosphingolipid levels has no impact on atherosclerosis in apolipoprotein E-null mice. J. Lipid Res. 49, 1677–1681 (2008); 10.1194/jlr.E800005-JLR200.

43. Voisine, M. et al. Age, sex, and valve phenotype differences in fibro-calcific remodeling of calcified aortic valve. J. Am. Heart Assoc. 9, e015610 (2020); 10.1161/JAHA.119.015610.

44. Summerhill, V. I., Moschetta, D., Orekhov, A. N., Poggio, P. & Myasoedova, V. A. Sex-specific features of calcific aortic valve disease. Int. J. Mol. Sci. 21; 10.3390(2020)/ijms21165620.

45. Büttner, P. et al. Dissecting calcific aortic valve disease-the role, etiology, and drivers of valvular fibrosis. Front. Cardiovasc. Med. 8, 660797 (2021); 10.3389/fcvm.2021.660797.

46. Nsaibia, M. J. et al. Implication of lipids in calcified aortic valve pathogenesis: why did statins fail? J. Clin. Med. 11 (2022); 10.3390/jcm11123331.

47. Schlotter, F. et al. Spatiotemporal multi-omics mapping generates a molecular atlas of the aortic valve and reveals networks driving disease. Circulation 138, 377–393 (2018); 10.1161/CIRCULATIONAHA.117.032291.

48. Laviad, E. L. et al. Characterization of ceramide synthase 2: tissue distribution, substrate specificity, and inhibition by sphingosine 1-phosphate. J. Biol. Chem. 283, 5677–5684 (2008); 10.1074/jbc.M707386200.

49. Nsaibia, M. J. et al. OxLDL-derived lysophosphatidic acid promotes the progression of aortic valve stenosis through a LPAR1-RhoA-NF-κB pathway. Cardiovasc. Res. 113, 1351–1363 (2017); 10.1093/cvr/cvx089.

50. Liu, S. et al. Plasma ceramides predict all-cause and cause-specific mortality in individuals with type 2 diabetes. J. Clin. Endocrinol. Metab. (2024); 10.1210/clinem/dgae388.

51. Hilvo, M. et al. Development and validation of a ceramide- and phospholipid-based cardiovascular risk estimation score for coronary artery disease patients. Eur. Heart J. 41, 371–380 (2020); 10.1093/eurheartj/ehz387.

52. Meeusen, J. W. et al. Plasma ceramides. Arterioscler. Thromb. Vasc. Biol. 38, 1933–1939 (2018); 10.1161/ATVBAHA.118.311199.

53. Laaksonen, R. et al. Plasma ceramides predict cardiovascular death in patients with stable coronary artery disease and acute coronary syndromes beyond LDL-cholesterol. Eur. Heart J. 37, 1967–1976 (2016); 10.1093/eurheartj/ehw148.

54. Lahelma, M. et al. The human liver lipidome is significantly related to the lipid composition and aggregation susceptibility of low-density lipoprotein (LDL) particles. Atherosclerosis 363, 22–29 (2022); 10.1016/j.atherosclerosis.2022.11.018.

55. Öörni, K., Jauhiainen, M. & Kovanen, P. T. Why and how increased plasma ceramides predict future cardiovascular events? Atherosclerosis 314, 71–73 (2020); 10.1016/j.atherosclerosis.2020.09.030.

56. Chang, Y. et al. Alcoholic and non-alcoholic fatty liver disease and associations with coronary artery calcification: evidence from the Kangbuk Samsung Health Study. Gut 68, 1667–1675 (2019); 10.1136/gutjnl-2018-317666.

57. Zhu, R.-R. et al. Non-alcoholic fatty liver disease is associated with aortic calcification: a cohort study with propensity score matching. Front. Endocrinol. 13, 880683 (2022); 10.3389/fendo.2022.880683.

58. Sinn, D. H. et al. Non-alcoholic fatty liver disease and progression of coronary artery calcium score: a retrospective cohort study. Gut 66, 323–329 (2017); 10.1136/gutjnl-2016-311854.

59. Schmidt, S. et al. Silencing of ceramide synthase 2 in hepatocytes modulates plasma ceramide biomarkers predictive of cardiovascular death. Mol. Ther. 30, 1661–1674 (2022); 10.1016/j.ymthe.2021.08.021.

60. Lenoir, G., D’Ambrosio, J. M., Dieudonné, T. & Čopič, A. Transport pathways that contribute to the cellular distribution of phosphatidylserine. Front. Cell Dev. Biol. 9, 737907 (2021); 10.3389/fcell.2021.737907.

61. Trinh, M. N. et al. Last step in the path of LDL cholesterol from lysosome to plasma membrane to ER is governed by phosphatidylserine. Proc. Natl. Acad. Sci. USA 117, 18521–18529 (2020); 10.1073/pnas.2010682117.

62. Boskey, A. L., Ullrich, W., Spevak, L. & Gilder, H. Persistence of complexed acidic phospholipids in rapidly mineralizing tissues is due to affinity for mineral and resistance to hydrolytic attack: in vitro data. Calcif. Tissue Int. 58, 45–51 (1996); 10.1007/BF02509545.

63. Tabassum, R., Widén, E. & Ripatti, S. Effect of biological sex on human circulating lipidome: An overview of the literature. Atherosclerosis 384, 117274 (2023); 10.1016/j.atherosclerosis.2023.117274.

64. Tabassum, R. et al. Lipidome- and genome-wide study to understand sex differences in circulatory lipids. J. Am. Heart Assoc. 11, e027103 (2022); 10.1161/JAHA.122.027103.

65. Holven, K. B. & van Roeters Lennep, J. Sex differences in lipids: A life course approach. Atherosclerosis 384, 117270 (2023); 10.1016/j.atherosclerosis.2023.117270.

66. Ishay, Y., Nachman, D., Khoury, T. & Ilan, Y. The role of the sphingolipid pathway in liver fibrosis: an emerging new potential target for novel therapies. Am. J. Physiol. 318, C1055–C1064 (2020); 10.1152/ajpcell.00003.2020.

67. Huwiler, A. & Pfeilschifter, J. Sphingolipid signaling in renal fibrosis. Matrix Biol. 68-69, 230–247 (2018); 10.1016/j.matbio.2018.01.006.

68. Ji, R. et al. Increased de novo ceramide synthesis and accumulation in failing myocardium. JCI Insight 2 (2017); 10.1172/jci.insight.82922 (2017).

69. Luft, F. C. Cystic fibrosis: the conductance regulator, ceramides, and possible treatments. J. Mol. Med. (Berl). 95, 1017–1019 (2017); 10.1007/s00109-017-1577-6.

70. Sato, M. et al. Modulation of transforming growth factor-beta (TGF-beta) signaling by endogenous sphingolipid mediators. J. Biol. Chem. 278, 9276–9282 (2003); 10.1074/jbc.M211529200.

71. Emma Le Nezet et al. Abstract 11668: Transcriptomic analysis reveals sex differences in gene expression profiling of stenotic aortic valves. Circulation 148, A11668–A11668 (2023); 10.1161/circ.148.suppl_1.11668.

72. Shah, T. A. & Rogers, M. B. Unanswered questions regarding sex and BMP/TGF-β signaling. J. Dev. Biol. 6 (2018); 10.3390/jdb6020014.

73. Kusminski, C. M. & Scherer, P. E. Lowering ceramides to overcome diabetes. Science 365, 319–320 (2019); 10.1126/science.aax6594.

74. Zhu, Q. & Scherer, P. E. Ceramides and atherosclerotic cardiovascular disease: a current perspective. Circulation 149, 1624–1626 (2024); 10.1161/CIRCULATIONAHA.123.065409.

75. Poss, A. M. et al. Machine learning reveals serum sphingolipids as cholesterol-independent biomarkers of coronary artery disease. J. Clin. Invest. 130, 1363–1376 (2020); 10.1172/JCI131838.

76. Leiherer, A. et al. Coronary event risk test (CERT) as a risk predictor for the 10-year clinical outcome of patients with peripheral artery disease. J. Clin. Med. 12 (2023); 10.3390/jcm12196151.

77. Folch, J., Lees, M. & Sloane Stanley, G. H. A simple method for the isolation and purification of total lipides from animal tissues. J. Biol. Chem. 226, 497–509 (1957).

78. Ni, Z., Angelidou, G., Lange, M., Hoffmann, R. & Fedorova, M. LipidHunter identifies phospholipids by high-throughput processing of LC-MS and shotgun lipidomics datasets. Anal. Chem. 89, 8800–8807 (2017); 10.1021/acs.analchem.7b01126.

79. Adams, K. J. et al. Skyline for Small Molecules: A unifying software package for quantitative metabolomics. J. Proteome Res. 19, 1447–1458 (2020); 10.1021/acs.jproteome.9b00640.

80. Lange, M. & Fedorova, M. Evaluation of lipid quantification accuracy using HILIC and RPLC MS on the example of NIST® SRM® 1950 metabolites in human plasma. Anal. Bioanal. Chem. 412, 3573–3584 (2020); 10.1007/s00216-020-02576-x.

81. Pang, Z. et al. MetaboAnalyst 6.0: towards a unified platform for metabolomics data processing, analysis and interpretation. Nucleic Acids Res. 52, W398–W406 (2024); 10.1093/nar/gkae253.

82. Burla, B. et al. MS-based lipidomics of human blood plasma: a community-initiated position paper to develop accepted guidelines. J. Lipid Res. 59, 2001–2017 (2018); 10.1194/jlr.S087163.

83. Kretschmer, F., Seipp, J., Ludwig, M., Klau, G. W. & Böcker, S. Small molecule machine learning: All models are wrong, some may not even be useful. Preprint at https://www.biorxiv.org/content/10.1101/2023.03.27.534311v2 (2024).

