## Supplementary figures and images for "Deep lipidomic profiling reveals sex dimorphism of lipid metabolism in fibro-calcific aortic valve disease"

### Extended data Figure 1

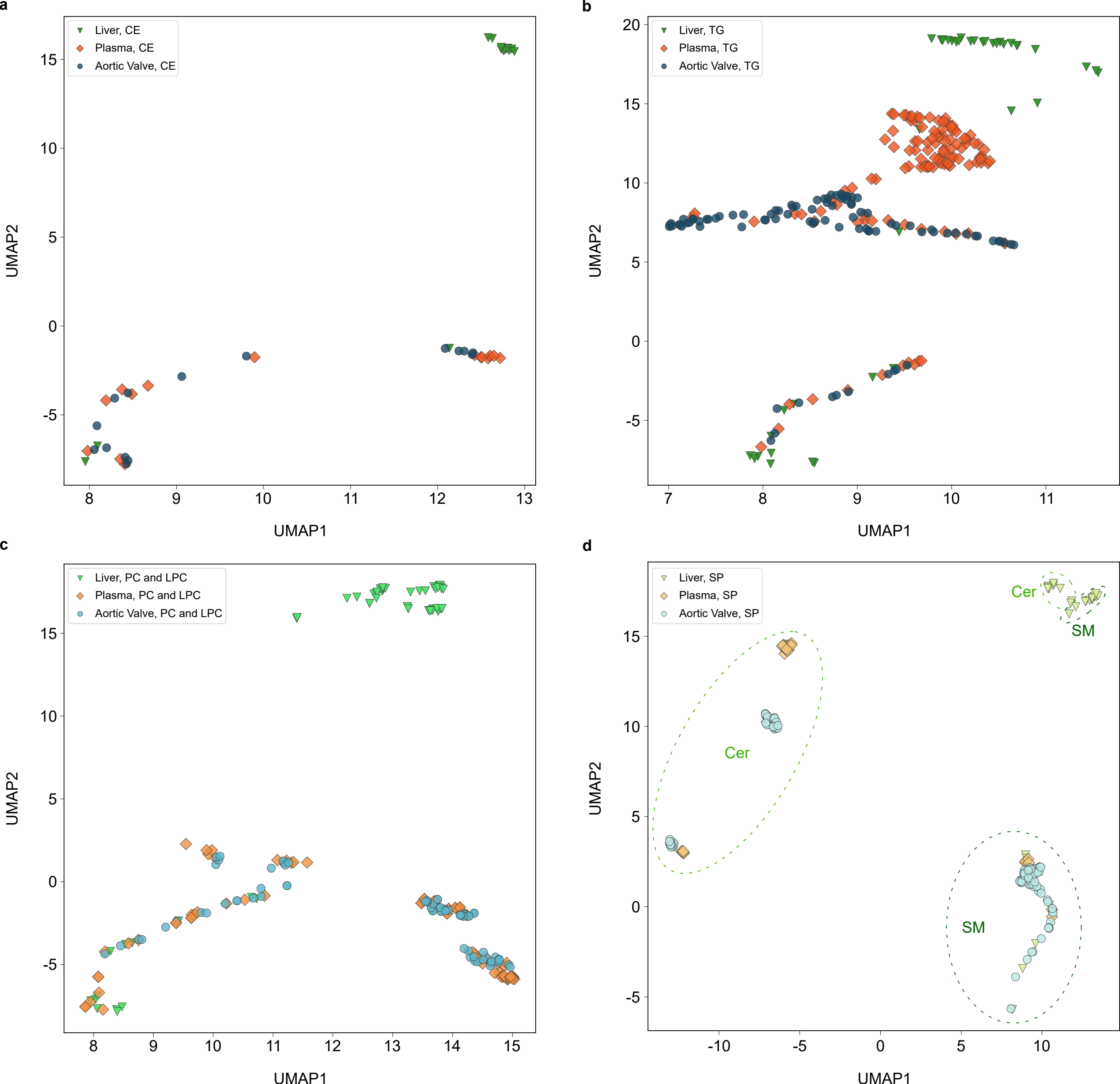

### Extended data Figure 2

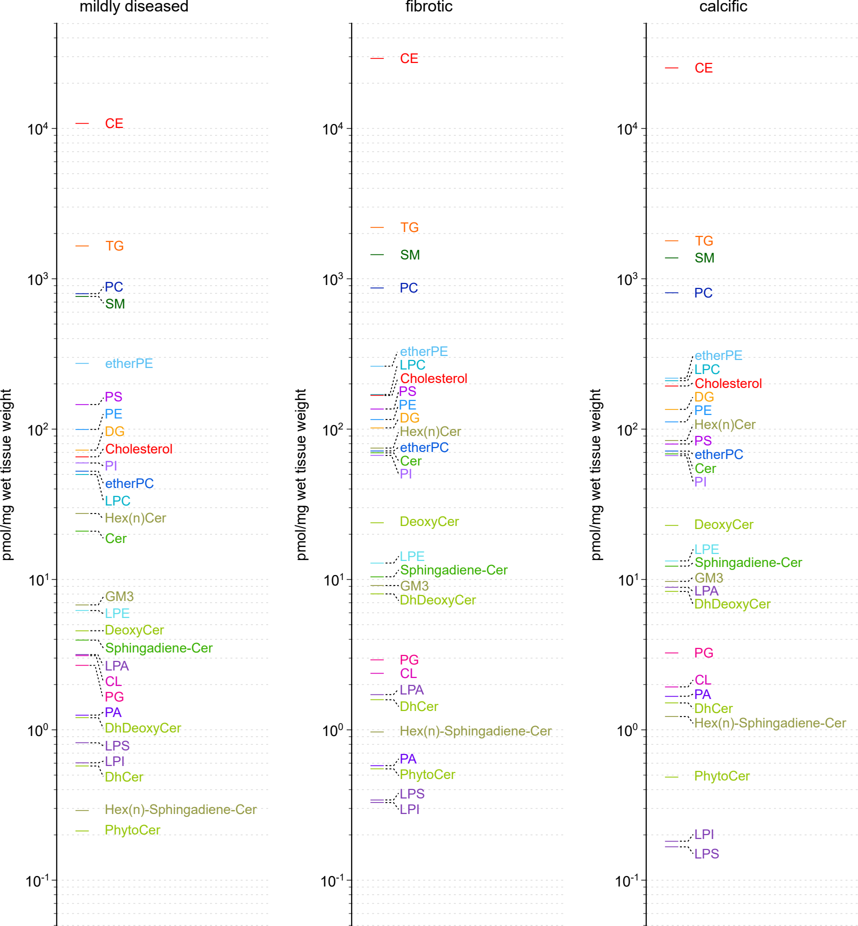

### Extended data Figure 3

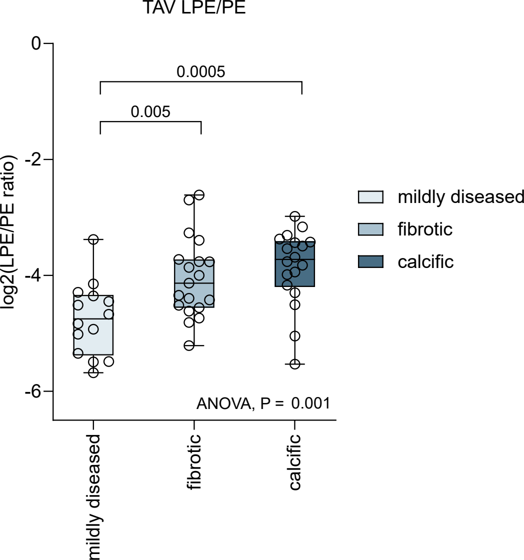

### Extended data Figure 4

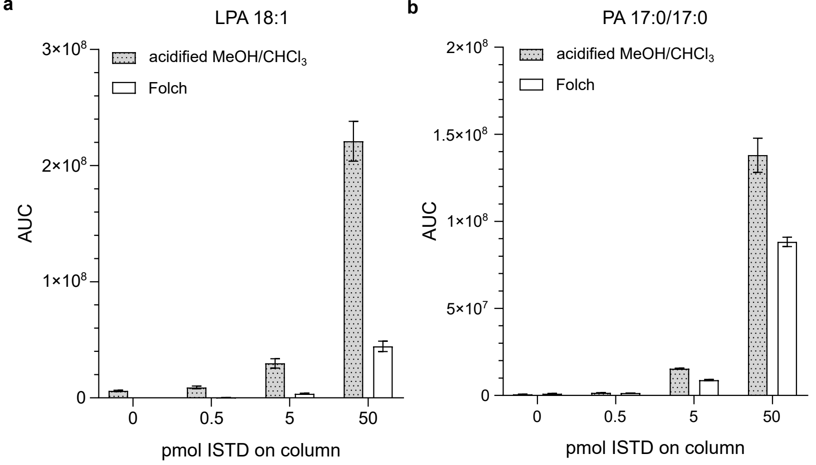


**c**

**Recovery**

| Extraction | pmol on column | LPA 18:1 | PA 17:0_17:0 |
|---|---|---|---|
| acidified $MeOH/CHCl_3$ | 0.5 | 90 ± 12 | 176 ± 14 |
| | 5 | 78 ± 11 | 169 ± 3 |
| | 50 | 61 ± 5 | 174 ± 12 |
| Folch | 0.5 | 11 ± 2 | 47 ± 2 |
| | 5 | 11 ± 2 | 54 ± 2 |
| | 50 | 16 ± 2 | 92 ± 3 |

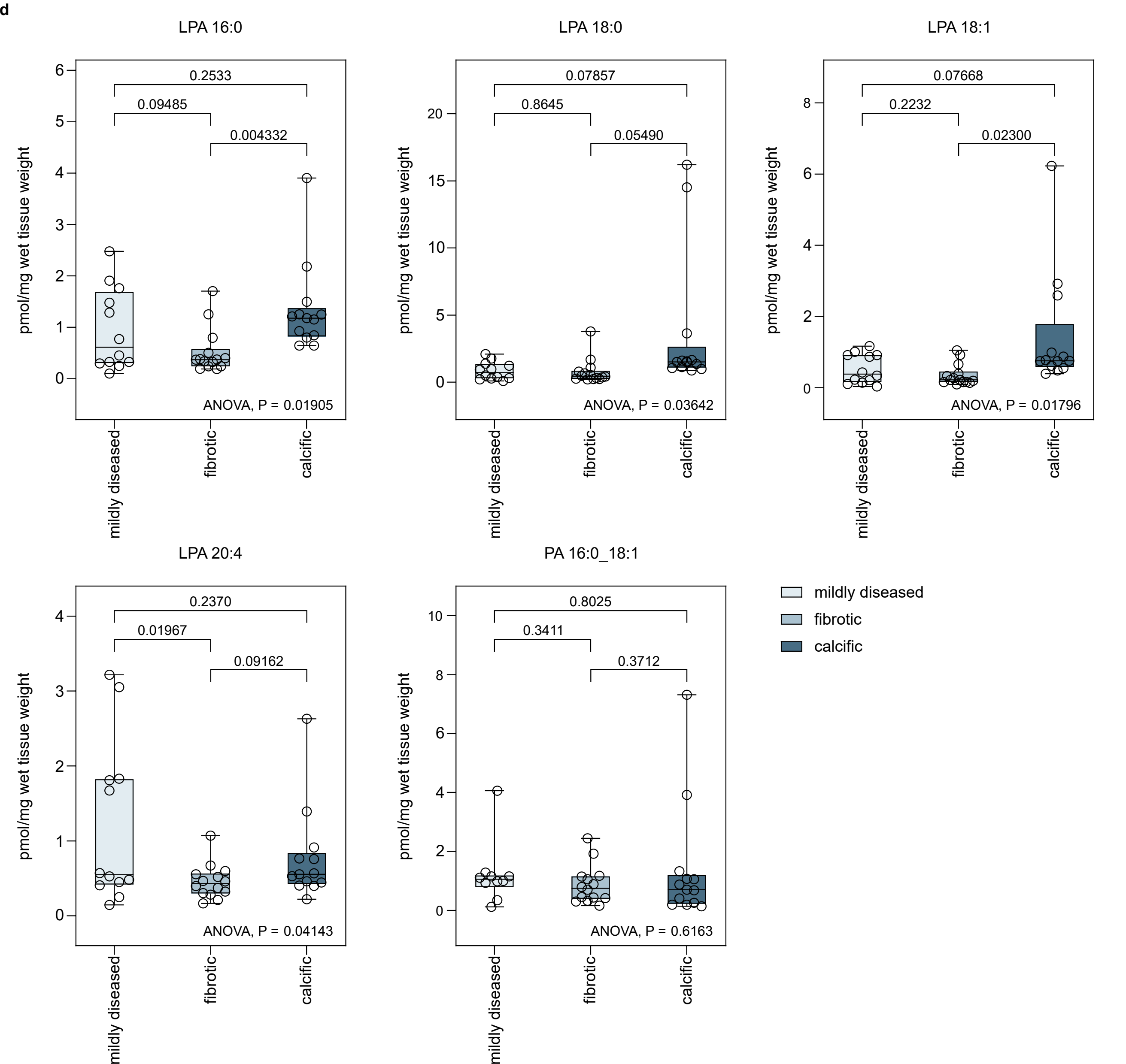
