## Supplementary Data Figures for "Deep lipidomic profiling reveals sex dimorphism of lipid metabolism in fibro-calcific aortic valve disease"

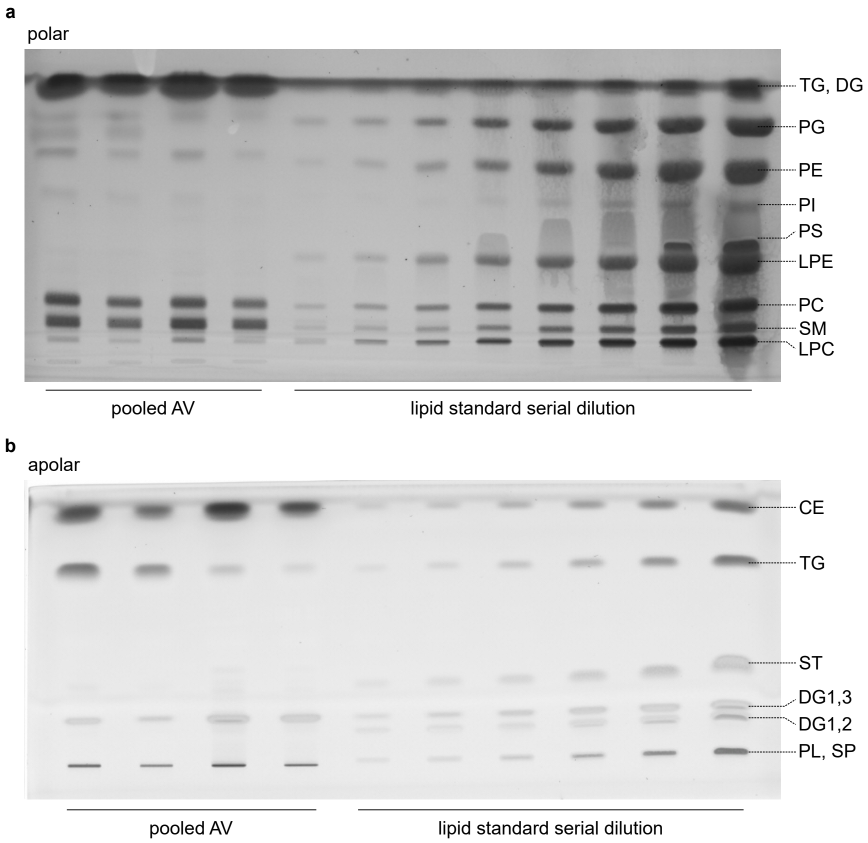


**Supplementary Data Fig. 1 | Quantitative high-performance thin-layer chromatograms (qHPTLC) for lipid class specific quantification of aortic valve lipids. a,** The polar plate was developed using chloroform/ethanol/trimethylamine/water (5:5:5:1, v/v/v/v). **b,** The apolar plate was developed using hexane/diethylether/acetic acid (85:15:1, v/v/v/v).

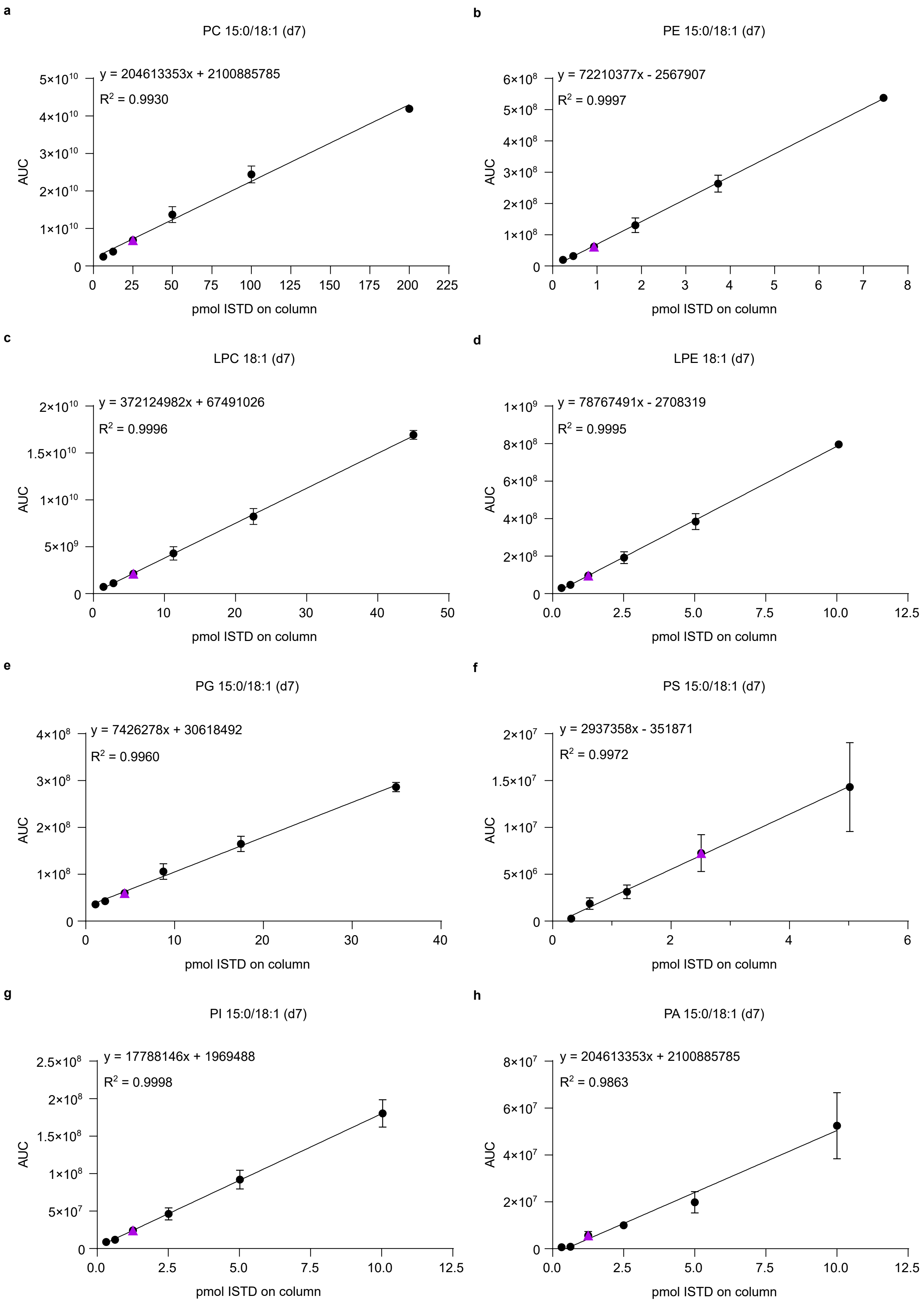
a
PC 15:0/18:1 (d7)
y = 204613353x + 2100885785
R2 = 0.9930
AUC
pmol ISTD on column
b
PE 15:0/18:1 (d7)
y = 72210377x - 2567907
R2 = 0.9997
AUC
pmol ISTD on column
c
LPC 18:1 (d7)
y = 372124982x + 67491026
R2 = 0.9996
AUC
pmol ISTD on column
d
LPE 18:1 (d7)
y = 78767491x - 2708319
R2 = 0.9995
AUC
pmol ISTD on column
e
PG 15:0/18:1 (d7)
y = 7426278x + 30618492
R2 = 0.9960
AUC
pmol ISTD on column
f
PS 15:0/18:1 (d7)
y = 2937358x - 351871
R2 = 0.9972
AUC
pmol ISTD on column
g
PI 15:0/18:1 (d7)
y = 17788146x + 1969488
R2 = 0.9998
AUC
pmol ISTD on column
h
PA 15:0/18:1 (d7)
y = 204613353x + 2100885785
R2 = 0.9863
AUC
pmol ISTD on column

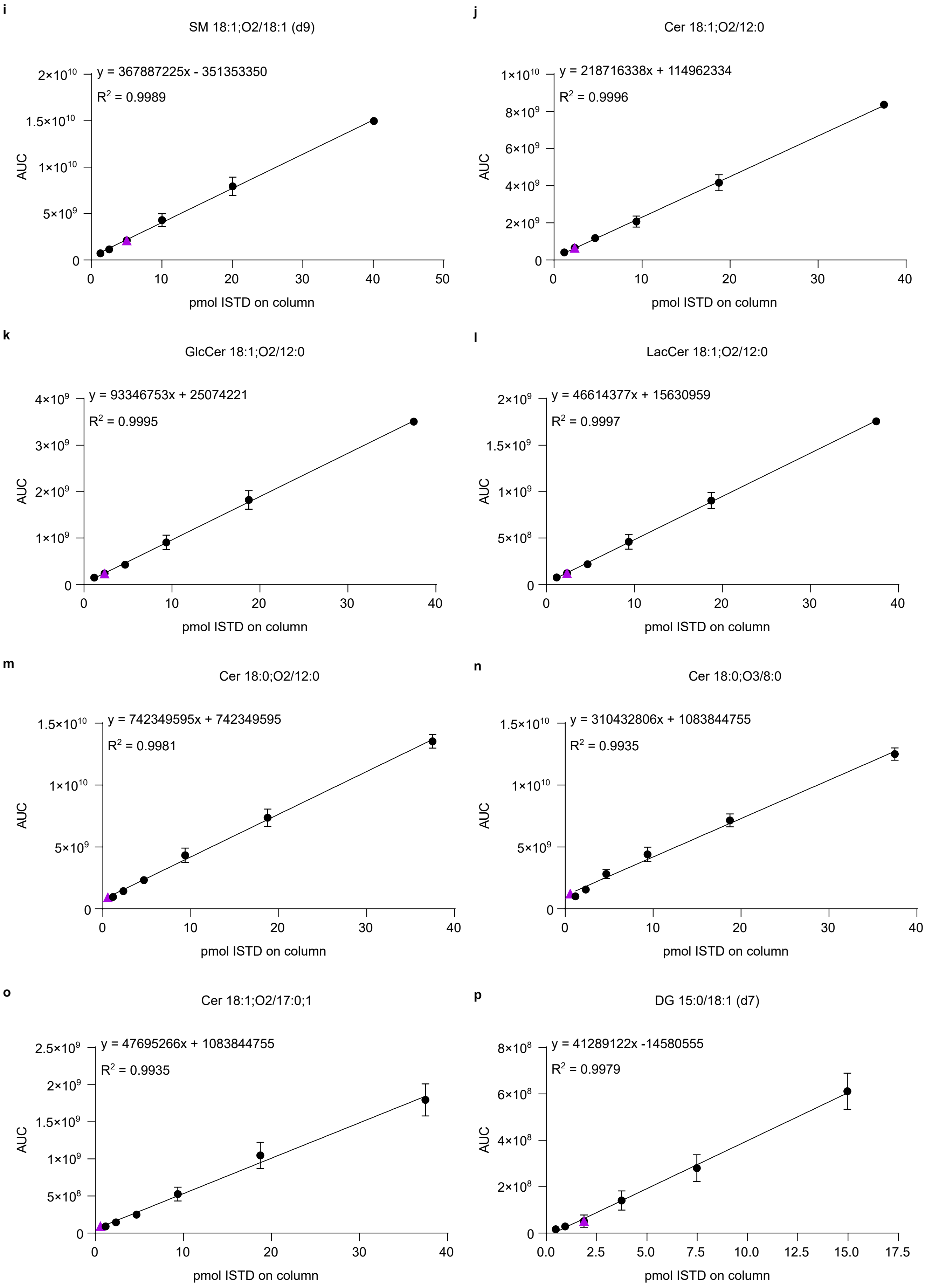

i
SM 18:1;O2/18:1 (d9)
y = 367887225x - 351353350
R² = 0.9989
AUC
pmol ISTD on column
j
Cer 18:1;O2/12:0
y = 218716338x + 114962334
R² = 0.9996
AUC
pmol ISTD on column
k
GlcCer 18:1;O2/12:0
y = 93346753x + 25074221
R² = 0.9995
AUC
pmol ISTD on column
l
LacCer 18:1;O2/12:0
y = 46614377x + 15630959
R² = 0.9997
AUC
pmol ISTD on column
m
Cer 18:0;O2/12:0
y = 742349595x + 742349595
R² = 0.9981
AUC
pmol ISTD on column
n
Cer 18:0;O3/8:0
y = 310432806x + 1083844755
R² = 0.9935
AUC
pmol ISTD on column
o
Cer 18:1;O2/17:0;1
y = 47695266x + 1083844755
R² = 0.9935
AUC
pmol ISTD on column
p
DG 15:0/18:1 (d7)
y = 41289122x -14580555
R² = 0.9979
AUC
pmol ISTD on column

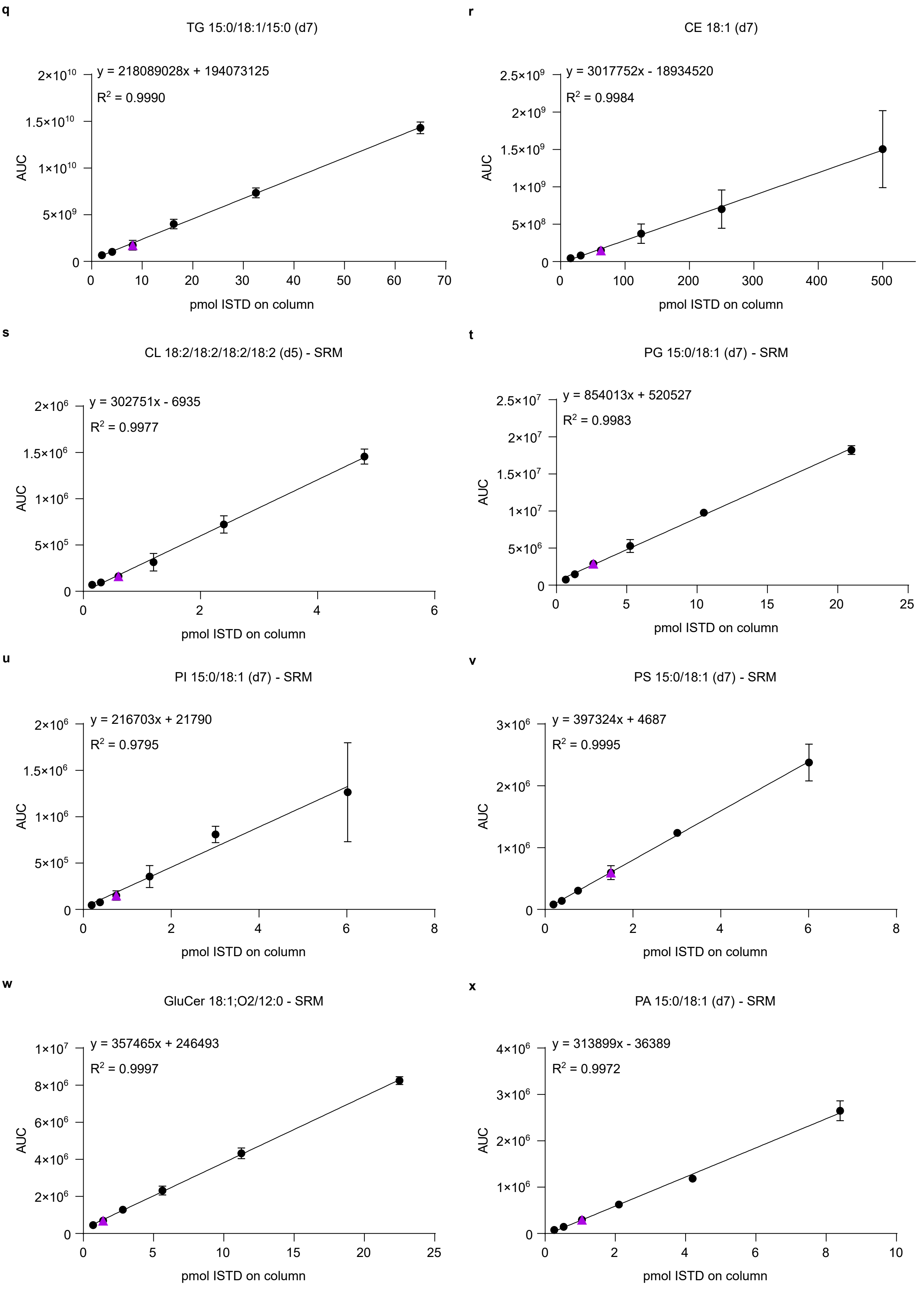
q
TG 15:0/18:1/15:0 (d7)
y = 218089028x + 194073125
R2 = 0.9990
AUC
pmol ISTD on column
r
CE 18:1 (d7)
y = 3017752x - 18934520
R2 = 0.9984
AUC
pmol ISTD on column
s
CL 18:2/18:2/18:2/18:2 (d5) - SRM
y = 302751x - 6935
R2 = 0.9977
AUC
pmol ISTD on column
t
PG 15:0/18:1 (d7) - SRM
y = 854013x + 520527
R2 = 0.9983
AUC
pmol ISTD on column
u
PI 15:0/18:1 (d7) - SRM
y = 216703x + 21790
R2 = 0.9795
AUC
pmol ISTD on column
v
PS 15:0/18:1 (d7) - SRM
y = 397324x + 4687
R2 = 0.9995
AUC
pmol ISTD on column
w
GluCer 18:1;O2/12:0 - SRM
y = 357465x + 246493
R2 = 0.9997
AUC
pmol ISTD on column
x
PA 15:0/18:1 (d7) - SRM
y = 313899x - 36389
R2 = 0.9972
AUC
pmol ISTD on column

**y**

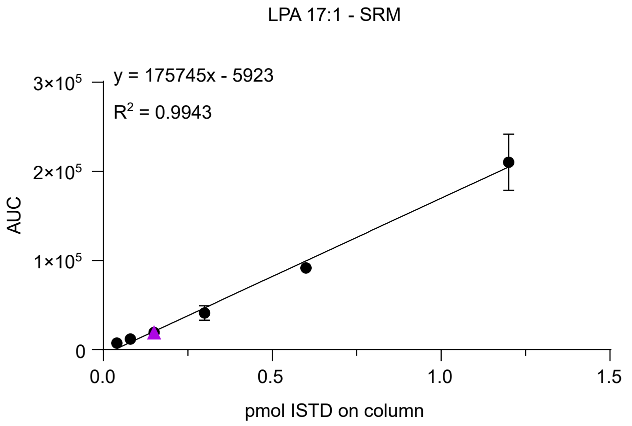


**z**

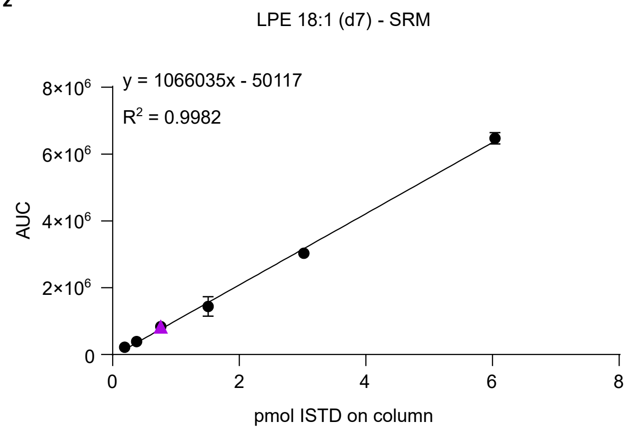


**Supplementary Data Fig. 2 | Area under curve (AUC) plotted against internal standard (ISTD) concentration in order to establish calibration curves for different AV lipid class standards. a - r,** Data used for the calibration curves was acquired using a Q Exactive Plus Hybrid Quadrupole Orbitrap mass spectrometer in full scan mode. **s – z,** Data used for the calibration curves was acquired using a TSQ Altis Plus Triple Quadrupole mass spectrometer in selected reaction monitoring (SRM)

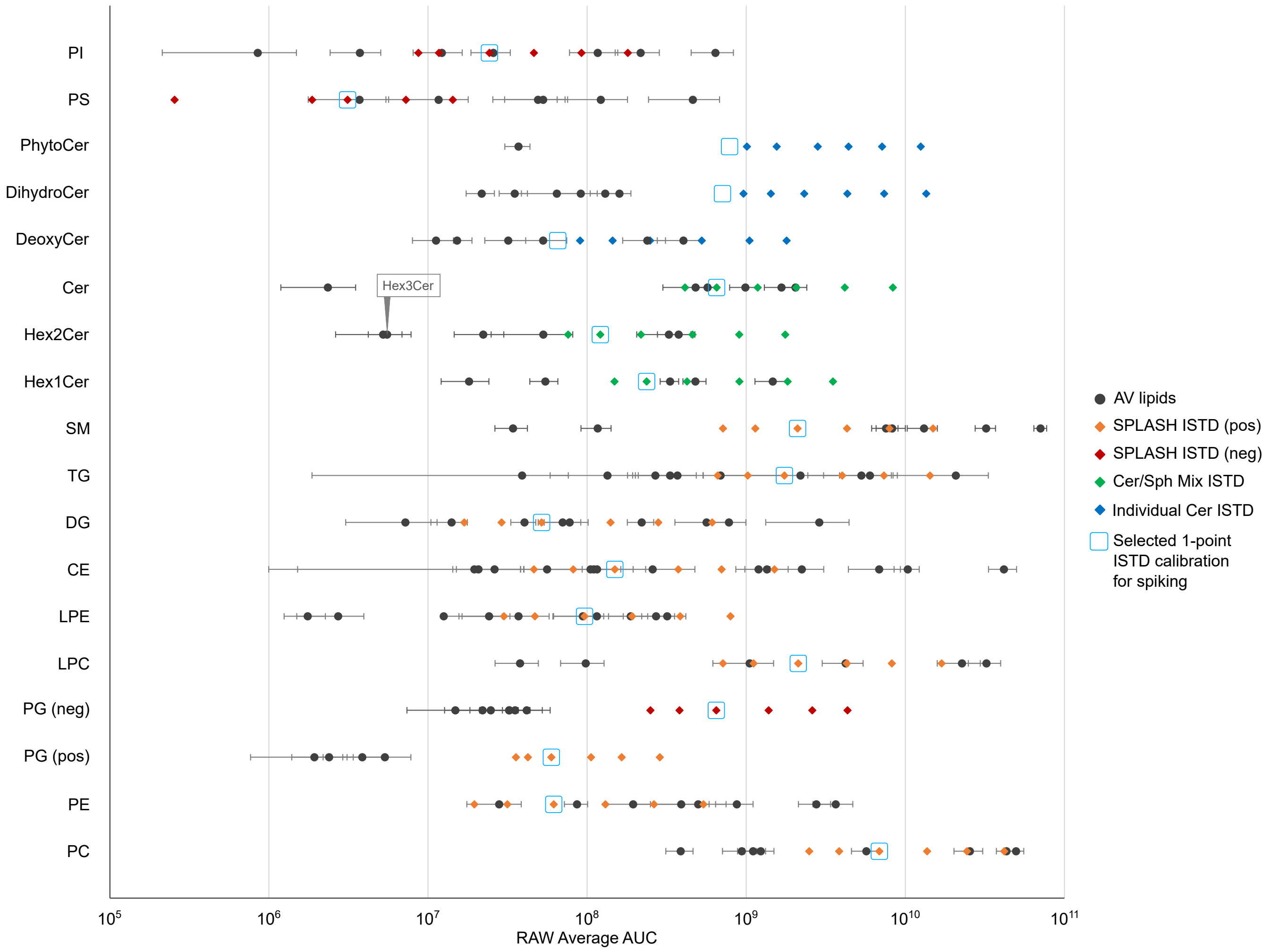


**Supplementary Data Fig. 3 | Concentration alignment of internal standard (ISTD) mixtures and endogenous aortic valve (AV) lipids.** Peak areas of five to nine endogenous lipids per subclass, representing some of the most, middle and the least abundant species of the class, were plotted alongside the peak areas of the corresponding lipid class ISTD in order to select the ideal ISTD amounts for each class (blue rectangle).
